# Combined linkage and association mapping of putative QTLs controlling black tea quality and drought tolerance traits

**DOI:** 10.1101/458596

**Authors:** Robert. K. Koech, Richard Mose, Samson M. Kamunya, Zeno Apostolides

## Abstract

The advancements in genotyping have opened new approaches for identification and precise mapping of Quantitative Trait Loci (QTLs) in plants, particularly by combining linkage and association mapping (AM) analysis. In this study, a combination of linkage and the AM approach was used to identify and authenticate putative QTLs associated with black tea quality traits and percent relative water content (%RWC). The population structure analysis clustered two parents and their respective 261 F1 progenies from the two reciprocal crosses into two clusters with 141 tea accessions in cluster one and 122 tea accessions in cluster two. The two clusters were of mixed origin with tea accessions in population TRFK St. 504 clustering together with tea accessions in population TRFK St. 524. A total of 71 putative QTLs linked to black tea quality traits and %RWC were detected in interval mapping (IM) method and were used as cofactors in multiple QTL model (MQM) mapping where 46 putative QTLs were detected. The phenotypic variance for each QTL ranged from 2.8–23.3% in IM and 4.1–23% in MQM mapping. Using Q-model and Q+K-model in AM, a total of 49 DArTseq markers were associated with 16 phenotypic traits. Significant marker-trait association in AM were similar to those obtained in IM, and MQM mapping except for six more putative QTLs detected in AM which are involved in biosynthesis of secondary metabolites, carbon fixation and abiotic stress. The combined linkage and AM approach appears to have great potential to improve the selection of desirable traits in tea breeding.

## Introduction

Tea plant (*Camellia sinensis* (L.) O. Kuntze) is an important economic crop grown in 36 tropical and semi-tropical countries around the world for production of tea beverages. In Africa, tea is mainly grown in Kenya, Malawi, Uganda, Tanzania, Zimbabwe, Rwanda, South Africa, Burundi, and Mauritius, although some tea is also grown in Ethiopia, Nigeria and Cameroon (FAO 2015a). Globally, Kenya is ranked third in terms of annual volume production of black tea after China and India and therefore, tea is Kenya’s top foreign exchange earner and export commodity among agricultural produces (Kenya National Bureau of Statistics 2012). Tea being universally the most popular beverage, the need to develop tea cultivars with optimum potential for black or green tea quality has recently become the single most important breeding objective. Currently, Kenya produces cut, tear and curl (CTC) black tea as the only product, for which the prices have declined owing to a global glut. This calls for concerted action, the chief being the diversification and added value of tea products, not only to help Kenya maintain its position as a leading exporter but also to enhance foreign exchange earnings.

Tea breeding involves the creation of genetic variability through controlled hybridisation between selected progenitors and evaluation of the newly generated progeny. This is to identify promising individuals with desirable traits with the aim of improving tea productivity. However, conventional tea breeding requires at least 20–25 years (Chen et al. 2013), from individual selection to local adaptability testing and to the final release of new tea cultivars to the farmers, hence requires considerable investment in terms of land, labour, time and money. Therefore, precise identification and screening of genetic resources, the development of molecular breeding technologies and the deep understanding of the genetics of important agronomic traits should be given more attention. Currently, the tea breeding program is primarily focused on high-yielding tea, better quality processed tea, genetic resistance to biotic and abiotic stresses prevailing in different tea growing regions and the regional adaptability. However, these agronomic traits in tea plant are quantitative in nature and not amenable to easy manipulations in breeding programs without elaborate and long-term field testing (Wachira et al. 2001; Kamunya et al. 2004). Thus, tea breeding improvement process would be more enhanced with the application of reliable and stable selection tools that are not prone to the continually changing environment. For these reasons, the selection of breeding materials by the evaluation of DNA markers for important agronomic traits that are controlled by a few loci could be preferable method for marker-assisted selection (MAS) for many traits in many different populations (Chen et al. 2013).

Linkage mapping and association mapping (AM) and are two methods widely used for locating genes or QTL in plant breeding (Xu et al. 2017). Linkage mapping has been used extensively for identifying the genetic basis of quantitative traits in plants (Kulwal 2018). However, linkage mapping is usually limited by low polymorphism or small population size and few recombination events, which are considered to estimate the genetic distances between marker loci causative genomic regions for QTLs (Collard et al. 2005). However, AM also known as Linkage Disequilibrium mapping has extensively been used to circumvent the limitations posed by linkage mapping in plants in the last few years (Kraakman et al. 2004). Association mapping identifies QTL by examining the trait-marker associations has been used to overcome the limitations of linkage mapping (Kraakman et al. 2004). In linkage mapping, only two alleles at any given locus can be studied in bi-parental crosses and gives low mapping resolution (Flint-Garcia et al. 2003). Thus, AM generally offers higher resolution mapping due to a greater number of recombination events. However, AM has a lower power to detect the effect of rare alleles compared to linkage analysis which has a higher statistical power due to greater allele replication (Korte and Farlow 2013). Moreover, population structure may cause false positives in AM, but this has been overcome by using a mixed-model, which take both population structure (Q) and kinship (K) into account to reduce false positives (Yu et al. 2006). Although linkage mapping and AM each offer more advantages over the other, they are often applied in conjunction to validate the QTL identified, thus reducing spurious associations.

The integrated approach of linkage-AM has been used in other crop plants such as *Arabidopsis* (Sterken et al. 2012), wheat (Shi et al. 2017) and maize (Li et al. 2016) to dissect quantitatively inherited traits. In tree species, an integrated method of linkage mapping and AM to decipher the nature of genetic architecture of potential QTLs for growth traits has been reported in poplar hybrids (Du et al. 2016) and maritime pine (Bartholomé et al. 2016). In grapes, (Fournier-Level et al. 2010) a combined linkage mapping and AM has been used to study the genetic patterns of anthocyanin content, which is a determinant of berry colour, in grapes. Recently, AM has been used to study the genetic relationship between tea caffeine synthase gene (TCS1) and the caffeine content in tea plant and its related species (Jin et al. 2016). In this current study, we integrated the high QTL detection power of the linkage mapping with the high-resolution power of AM. The linkage-AM approach will not only accelerate the pace of QTL mapping in tea breeding improvement but will also precisely identify reliable QTLs linked to black tea quality and drought tolerance in tea. The objective of this study was to integrate linkage mapping and AM to precisely identify and authenticate putative QTLs linked to caffeine, catechins, theaflavins, tea tasters’ scores, and %RWC.

## Material and methods

### Population type

Two pseudo-testcross populations used in this current study consisted of 109 F_1_ progeny from TRFK St. 504 (TRFK 303/577 × GW Ejulu) and 152 F_1_ progeny from TRFK St. 524 (GW Ejulu × TRFK 303/577) as described previously (Koech et al. 2018). In brief, the two parental clones, TRFK 303/577 and GW Ejulu, are of Assamica and China varieties, respectively. The two clones were chosen on the basis of their contrasting attributes. GW Ejulu is a low-yielding, high black tea quality and moderate levels of caffeine, but high in total catechins and individual catechins contents. Clone TRFK 303/577, which is an open-pollinated progeny of clone TRFK 6/8, is high-yielding, drought-tolerant, medium in black tea quality, caffeine and individual catechins.

### Phenotypic data

A total of 16 phenotypic traits as described in the previous work by Koech et al. (2018) were used in this study. In brief, all the 16 phenotypic traits were assessed for a normal distribution. The mean and the standard deviation for each phenotypic trait in each parent and the progenies were calculated. The significance difference between parental values and progenies was analysed using Student’s t test.

### Population structure

The population structure was inferred from the 1,421 DArT markers data previously used to construct genetic linkage map using the STRUCTURE software version 2.3.4 (Pritchard et al. 2010). Twenty independent runs were carried out using the following parameters; number of populations (K) from 1 to 10, burn-in time and Markov Chain Monte Carlo (MCMC) replication number were both set to 100,000 for model of admixture with correlated allele frequencies. The natural logarithms of the probability data (LnP(K)) and the *ad hoc* delta K statistical were calculated using STRUCTURE Harvester (Earl 2012), and the optimal K according to the delta K value was then selected (Mezmouk et al. 2011). Finally, the population structure matrix (Q) was obtained by integrating 20 independent replicate runs and applying CLUMPP software (Jakobsson and Rosenberg 2007). STRUCTURE bar plots were plotted using STRUCTURE Plot v 2.0 (Ramasamy et al. 2014).

### Linkage mapping, QTL analyses and allelic effects

One thousand four hundred and twenty-one DArTseq markers derived from *C. sinensis* genomic DNA of the two parents and 261 F_1_ progeny were subjected to linkage mapping analyses using JoinMap 4.0 (Van Ooijen 2006) and QTL mapping analyses using MapQTL 6.0 software (Van Ooijen *et al.*, 2000) as described previously (Koech et al. 2018; Koech et al., 2019).

The allelic effects were estimated as A_f_□=□[(µac+µad)−(µbc+µbd)]/4 for female additivity, A_m_□=□[(µac+µbc)−(µad+µbd)]/4 for male additivity and D□=□[(µac+µbd)−(µad+µbc)]/4 for dominance where µac, µad, µbc and µbd are estimated phenotypic means associated to each of the four possible genotypic classes ac, bc, ad and bd, derived from ab×cd cross (Sadok et al, 2013).

### Association mapping

Using the Q and K matrices data files from STRUCTURE software as a covariates, 1,421 DArT markers of the 261 F_1_ progeny were tested for association with each phenotype using a general linear model (GLM), mixed linear model (MLM) that included kinship, phylogenetic tree in TASSEL (trait analysis by association, evolution and linkage) Version 5.2.43 software (Bradbury et al. 2007). Principal component analysis (PCA) was performed using JMP Pro 14 to visualise the dispersion of the association panel in a graph. The Q and kinship (K) matrices were used to correct the effects of population substructure in the association panel which can cause false-positive associations. The p-value and R^2^ were used to determine whether a QTL is associated with the marker and the magnitude of QTL effects, respectively.

## Results

### Frequency distribution

A wide range of variation in individual catechins, theaflavins, and caffeine contents was observed in the two parents and F_1_ progeny for all 16 phenotypic traits as described previously by Koech et al. (2018).

### Population structure

A population structure analysis showed that the value of Evanno’s ΔK presented a sharp spike at K = 2, which suggested that this population panel was clustered into two groups (Figure 1). The two groups or clusters are represented by two clades with parental clone TRFK 303/577 clustering with 141 progenies in cluster one (blue) and parental clone GW Ejulu (not shown) clustering with 122 progenies in cluster two (black), respectively (Figure 2). The two clusters were accession with mixed origin with tea accessions in TRFK St. 504 population clustering together with tea accessions in TRFK St. 524 population. Furthermore, the results from PCA (Figure 3) and a phylogenetic tree-based on Nei’s genetic distance was in agreement with the structure analysis results (Figure 4). The population structure of the two tea populations was examined with the neighbour-joining algorithm using Euclidian distance on the DArTseq marker band intensities.

**Figure 1.**
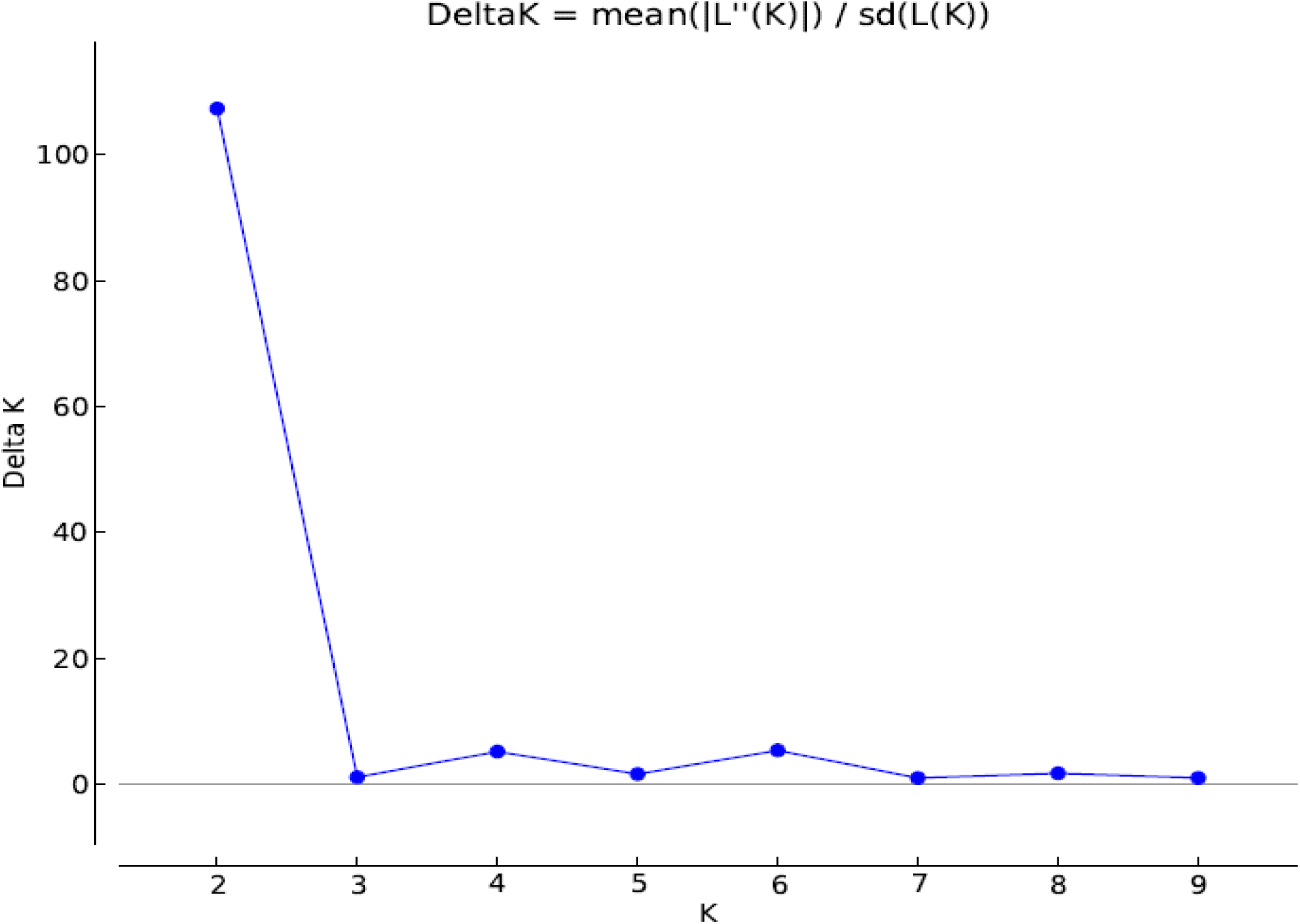
Delta K values plotted from 1 to 10 for 261 tea accessions panel.

**Figure 2.**
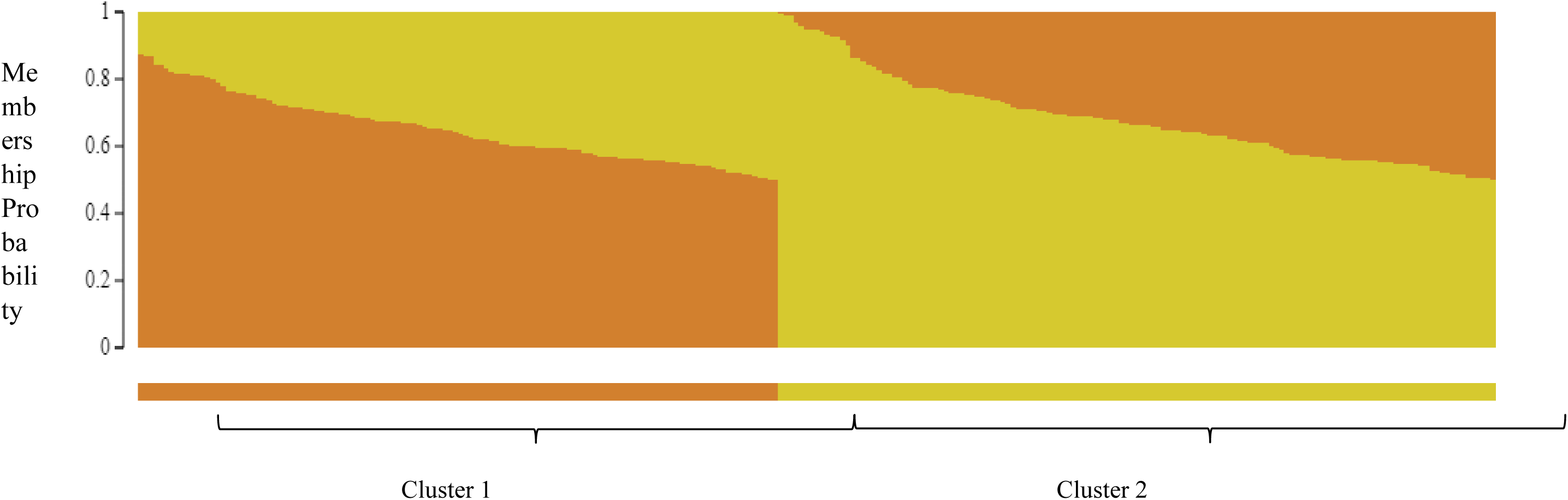
Population structure of 261 tea accessions panel (K = 2)

**Figure 3.**
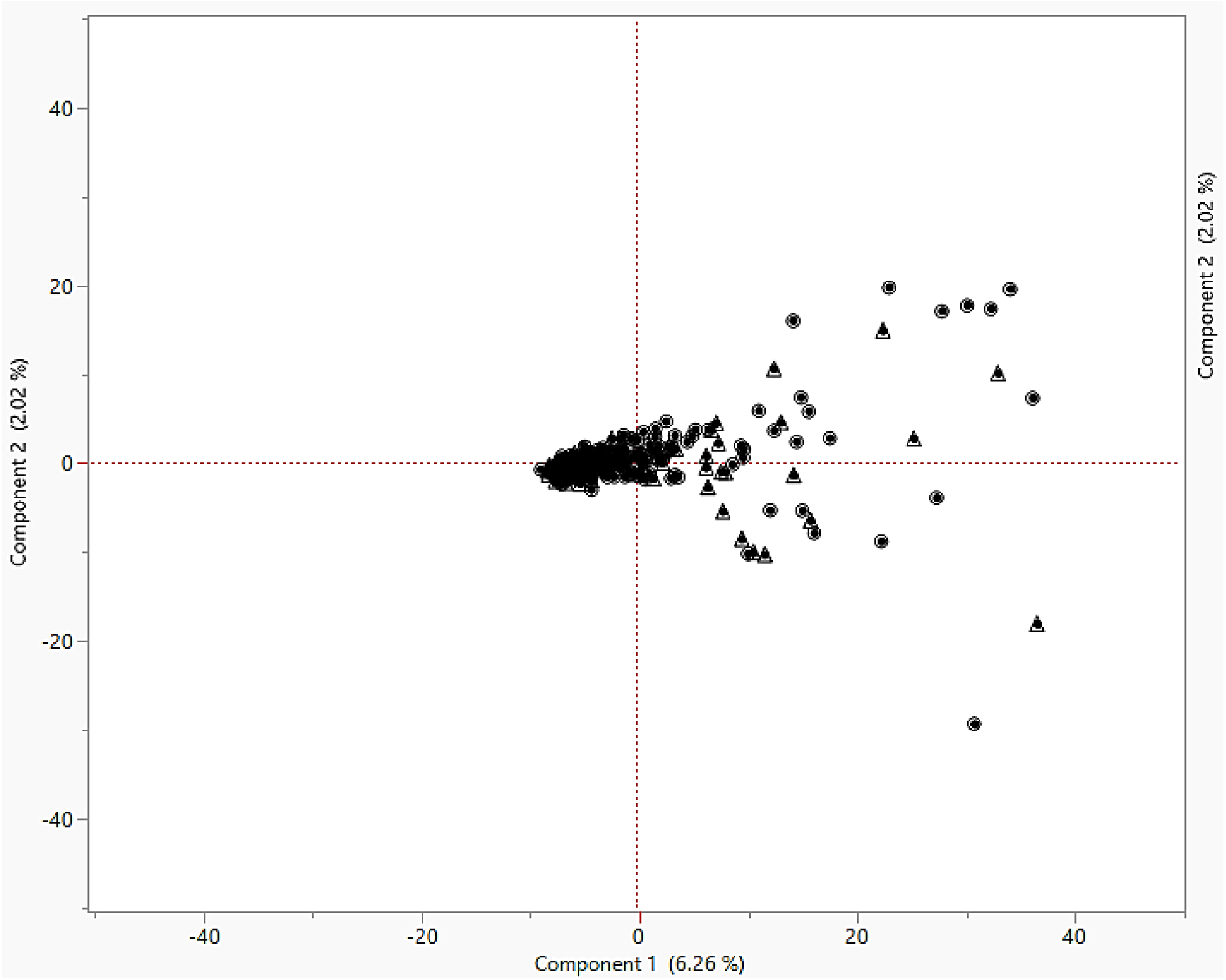
Principal component analysis of 261 tea accessions. 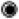 represents cluster 1, 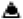 represents cluster 2

**Figure 4.**
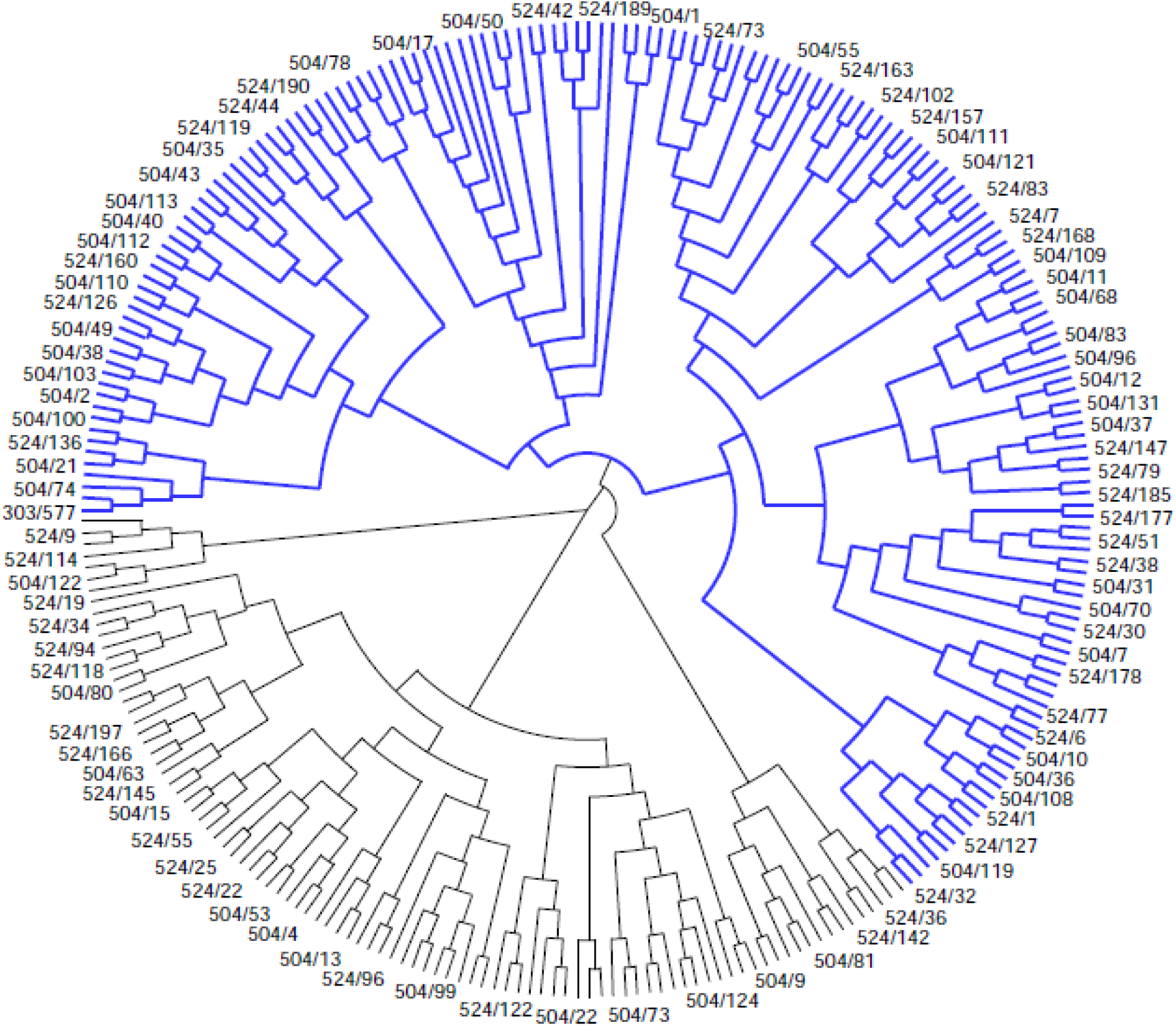
Phylogenetic tree-based on Nei’s genetic distance. Blue clade represents cluster 1, black clade represents cluster 2.

### Linkage mapping and association mapping

#### Interval mapping and allelic effects

The genetic linkage map was constructed using a total of 1,421 DArT markers as described previously (Koech et al. 2018). A total of 71 putative QTLs were identified for the 16 phenotypic traits measured using the interval mapping (Table 1). Of all the putative QTLs identified, most QTLs were found in almost all linkage groups, except LG03, LG05 and LG11. For black tea quality traits, 67 putative QTLs were detected and were located on almost all linkage groups, except on LG03, LG05 and LG11. The remaining three putative QTLs which are associated with drought tolerance were identified in LG02, LG06 and LG09. Eight putative QTLs were identified on LG01, seven in LG02 and LG12, 10 in LG04, five in LG06 and LG15, three in LG07, LG08, LG09, two in LG10, 15 in LG13, four in LG15, respectively. The highest number of putative QTLs were associated with catechin (13), EGC (12), tea tasters’ score (12), caffeine (10) and EGC (9), respectively. The phenotypic variance for all the identified putative QTLs ranged from 2.8% for EGC to 23.3% for qECG, respectively. The phenotypic variance for %RWC ranged from 5.7 to 7.3%, while for the tea tasters’ score, it ranged from 5.8 to 9.1%. Half of the allelic effects identified to be associated with black tea quality and drought tolerance traits had a positive additive effect. There were eight allelic effects associated with female additivity, and 18 were associated with male additive effects, respectively. The highest positive female and male additive effects were qEGC and qECG, respectively. Although, some putative QTLs also exhibited high negative allelic effects, which were from either female or male additive effects or both, such as qEGCG, qCAT, qEC and qCaffeine (Table 1). All the putative QTLs identified in IM were used as cofactors in MQM mapping.

**Table 1.**
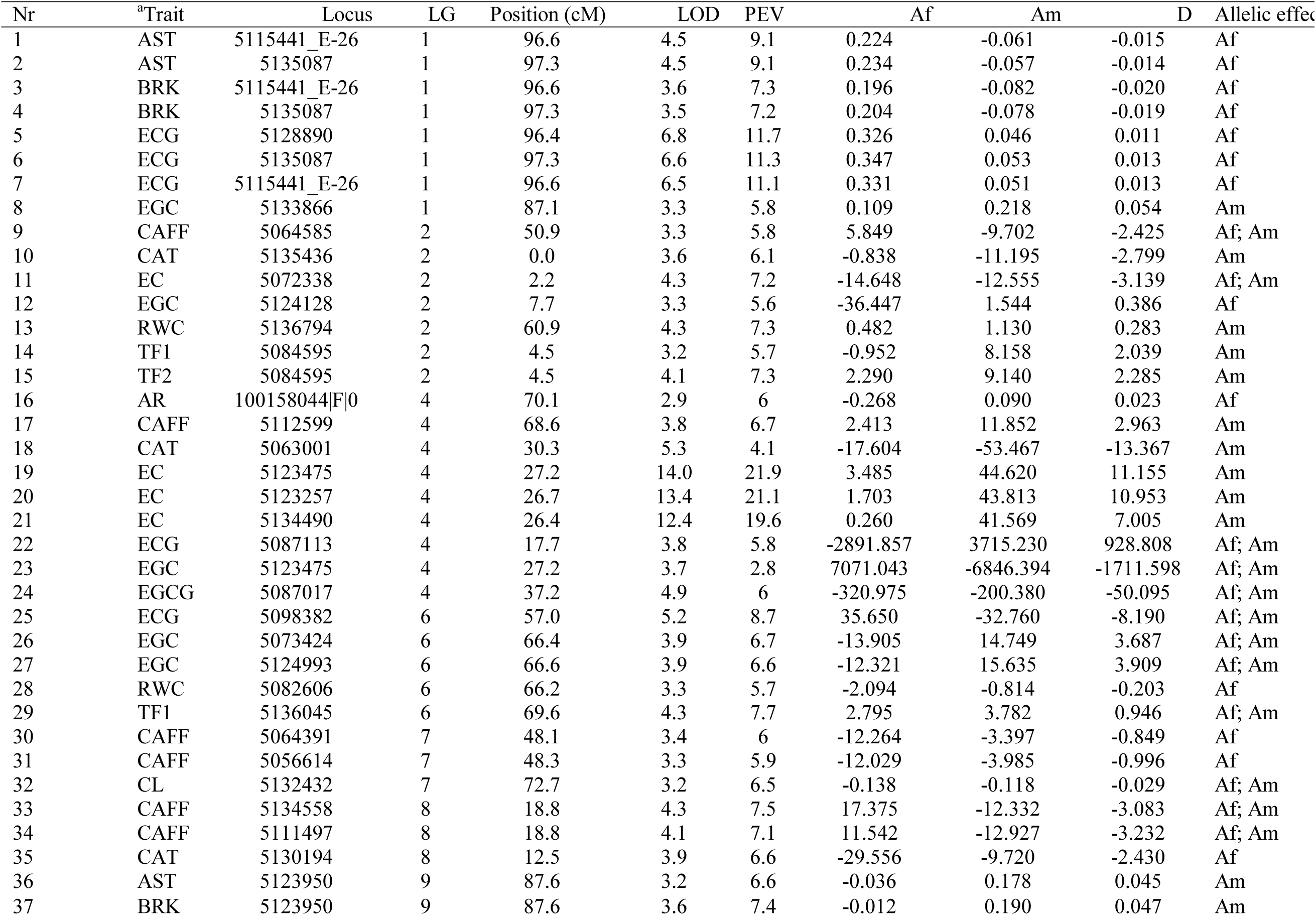
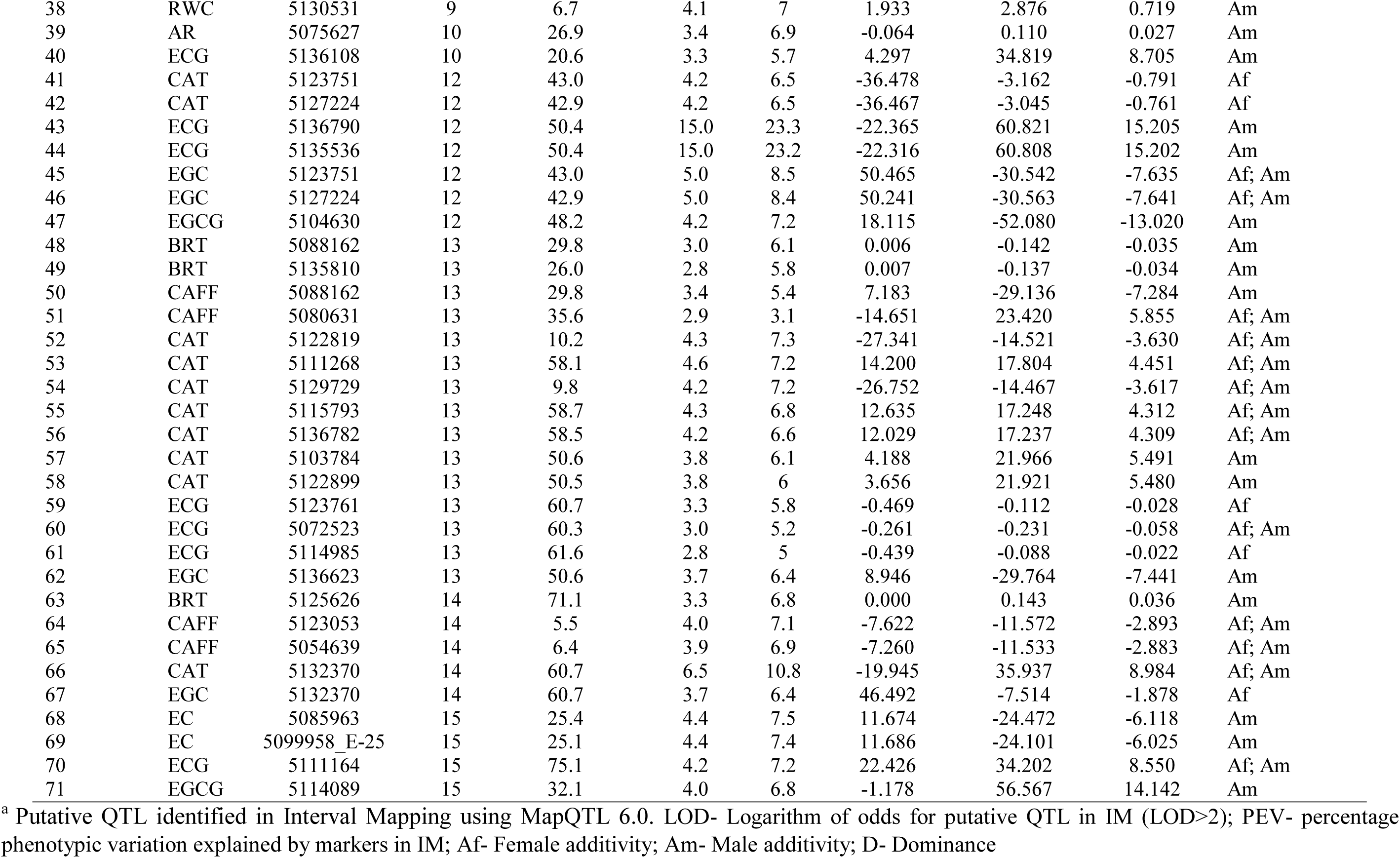
Linkage mapping results for putative QTLs in black tea quality and drought tolerance traits in St. 504 and St. 524 using interval mapping in MapQTL 6.0

#### Multiple QTL model mapping and allelic effects

A total of 46 putative QTLs (LOD > 3.0) were identified in 16 phenotypic traits using MQM mapping method from the selected cofactors in IM. Similarly, no putative QTLs in LG03, LG05 and LG11, respectively were associated with any of the phenotypic traits. Most of the putative QTLs previously identified in IM were also identified MQM mapping. Of these 46 putative QTLs identified, 43 putative QTLs that were associated with black tea quality traits while three were associated with qRWC (Table 2). The three putative QTLs associated with qRWC were similar to those identified in IM. Four putative QTLs were identified in LG01 and LG12, seven in LG02, five in LG04, three in LG06, LG09, LG13, LG14 and LG15, two in LG07 and LG08 and one in LG01, respectively. A similar trend was observed in the number of putative QTLs for each trait as in IM with a high putative QTL number for catechin, ECG, EGC and caffeine, respectively. The phenotypic variance for all the identified putative QTL ranged from 4.1% for qCAT to 23% for qECG, respectively. The phenotypic variance for qRWC was similar to IM results, and it ranged from 5.7 to 7.3%, while for the tea tasters’ score, it ranged from 6.0 to 9.1%. For allelic effects, similar results to IM with qEGC and qECG showing high positive female additive and male additive effects, respectively, this trend was also observed in MQM mapping. The putative QTLs identified using both IM, and MQM mapping were almost similar although the efficiency and the accuracy of detecting QTL is achieved by employing MQM instead of the single QTL model used in IM.

**Table 2.**
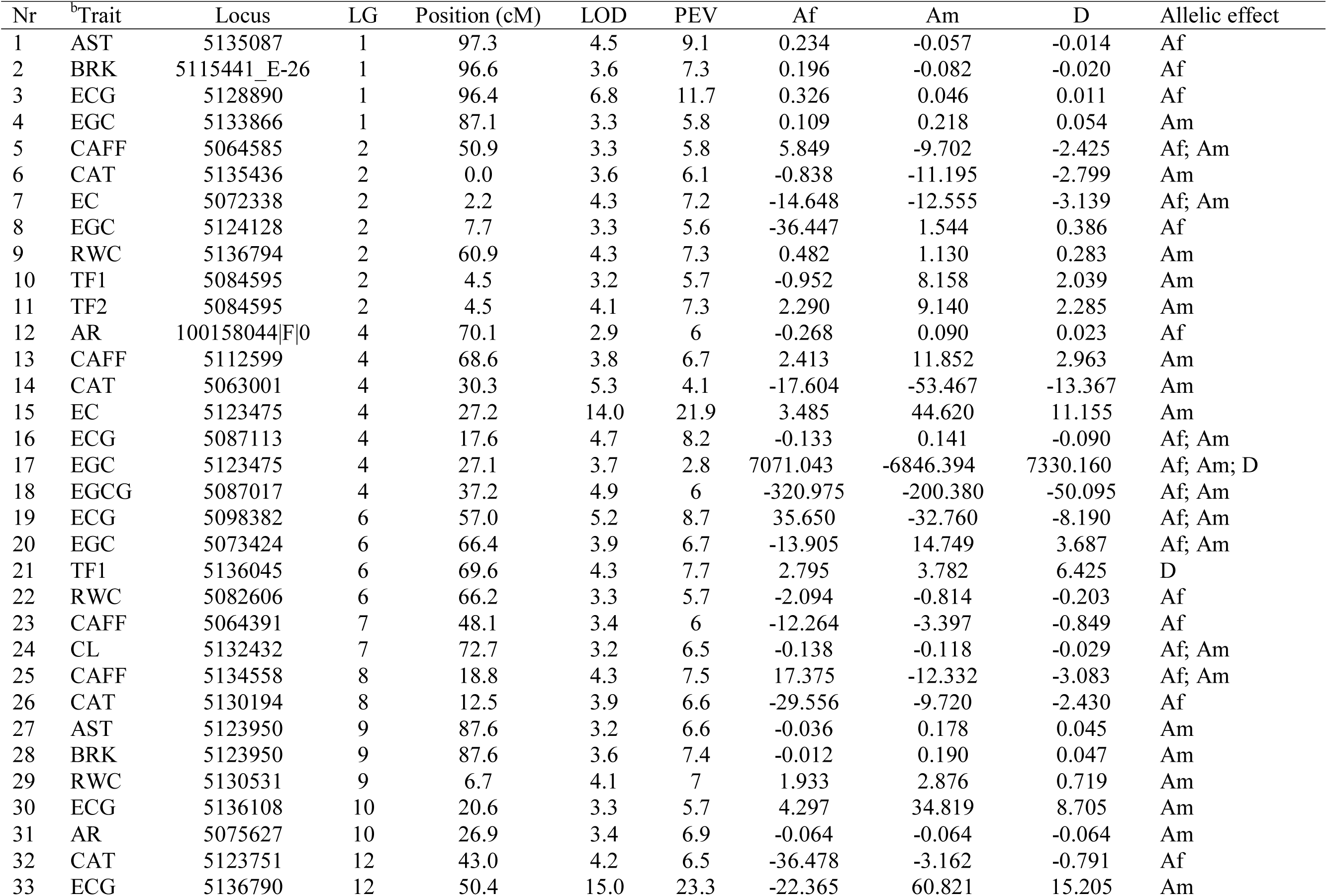
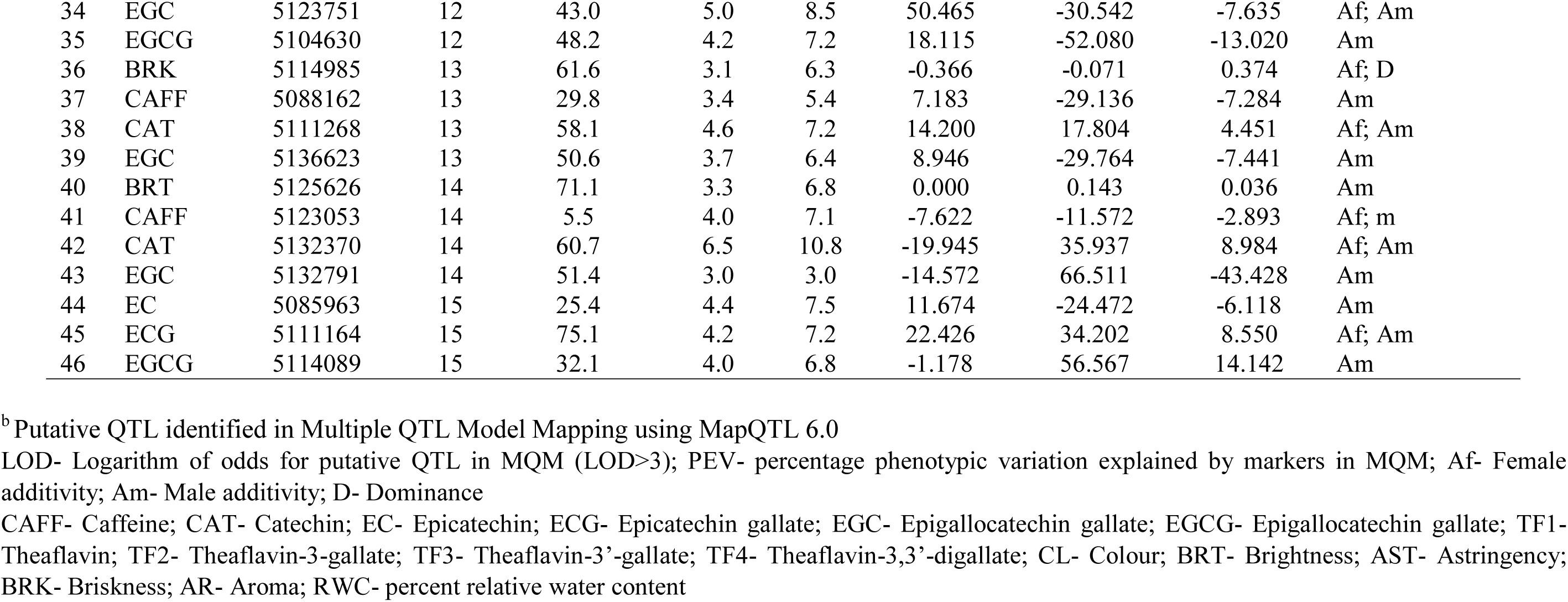
Linkage mapping for putative QTLs in black tea quality and drought tolerance traits in St. 504 and St. 524 using multiple QTL model mapping in MapQTL 6.0

#### Association mapping

Four models GLM, GLM (Q), mixed linear model with kinship matrix (MLM) (K) and mixed linear model with kinship matix and population structure matrix (MLM) (Q+K) in TASSEL software v5.2.43 were used to determine AM and the effects of population structure on the AM to reduce the inflation of false-positive associations. The p-values were plotted cumulatively for each model, and the distribution examined. However, no further analysis was done using GLM model since marker-trait association results were characterised by excess small p-values, which indicated an abundance of spurious associations (Supplementary Figure 1). The association analysis using the Q-model and Q+K-model detected a total of 49 DArT markers associated with 15 different black tea quality traits and one drought-tolerant trait. The GLM (Q) model showed associations between DArT markers and traits (p < 0.01) and was confirmed using the MLM (Q+K) model (Table 3, Supplementary Figures 2–4). The MLM Q+K-model (MLM with Q-matrix and K-matrix as a correction for population structure) showed a good fit for the p-values (p < 0.01), as compared to GLM (Q) models, which were characterised by a few associations with excess of small p-values, which indicates abundance of spurious associations (Table 3, Supplementary Figures 2 and 4). Also, the MLM Q+K-model also showed better small p-values than the K-model (MLM with K-matrix as a correction for population structure) (Supplementary Figures 3 and 4). The GLM (Q) model may not have accounted for the heterogeneity of the genetic background in some tea cultivars under study, which may have resulted in false-positive associations. Ideally, the distribution of p-values should follow a uniform distribution with less deviation from the expected p-values. However, the two models were used to compare the effect of population matrix and kinship matrix on GLM and MLM, respectively, in associating mapping. Therefore, taking into account the performance of the two different models, the results from the MLM (Q+K) model appeared to have controlled better population structure and kinship relationships than GLM (Q) model. While it might be tempting to consider p-values that remain extreme after MLM (Q+K) model correction as true associations, we need to keep in mind that minor allele frequency also influences p-values.

**Table 3.** Association mapping results for black tea quality and drought tolerance traits in St. 504 and St.524 using the GLM (Q) and MLM (Q+K) method, respectively (p <0.01)

| Nr | Trait | Marker | LG | Position (cM) | p-Value | Marker R<sup>2</sup> GLM (Q) | p-Value | Marker R<sup>2</sup> MLM (Q+K) |
| --- | --- | --- | --- | --- | --- | --- | --- | --- |
| 1 | ECG | 5128890 | 1 | 96.4 | 1.18E-03 | 0.042 | 5.31E-03 | 0.031 |
| 2 | EGC | 5086476 | 1 | 87.5 | 8.20E-03 | 0.031 | 8.54E-04 | 0.043 |
| 3 | EGC | 5122985 | 1 | 87.5 | 1.47E-04 | 0.056 | 2.41E-03 | 0.038 |
| 4 | EGC | 5133866 | 1 | 87.1 | 8.94E-03 | 0.030 | 6.62E-04 | 0.046 |
| 5 | CAFF | 5134416 | 2 | 44.4 | 2.46E-04 | 0.057 | 5.58E-04 | 0.050 |
| 6 | CAFF | 5136551 | 2 | 43.3 | 9.60E-04 | 0.048 | 7.51E-03 | 0.030 |
| 7 | CAFF | 5137282 | 2 | 50.7 | 2.27E-03 | 0.042 | 9.72E-03 | 0.026 |
| 8 | EC | 5132500 | 2 | 7.9 | 2.91E-03 | 0.036 | 1.37E-04 | 0.052 |
| 9 | RWC | 5136794 | 2 | 60.9 | 8.58E-03 | 0.032 | 3.02E-03 | 0.033 |
| 10 | CAT | 5063001 | 4 | 30.3 | 2.34E-09 | 0.098 | 6.67E-08 | 0.109 |
| 11 | CAT | 5064764 | 4 | 28.8 | 6.42E-37 | 0.326 | 7.80E-22 | 0.379 |
| 12 | CAT | 5136945 | 4 | 28.1 | 6.39E-40 | 0.345 | 1.81E-21 | 0.406 |
| 13 | EC | 5112438 | 4 | 27.4 | 4.90E-07 | 0.087 | 1.50E-05 | 0.067 |
| 14 | EC | 5123257 | 4 | 26.7 | 3.89E-08 | 0.101 | 7.16E-07 | 0.089 |
| 15 | EC | 5134490 | 4 | 26.4 | 5.15E-19 | 0.228 | 6.21E-15 | 0.246 |
| 16 | EC | 5136058 | 4 | 27.6 | 1.00E-06 | 0.083 | 5.31E-06 | 0.073 |
| 17 | EC | 5136410 | 4 | 29.9 | 2.39E-05 | 0.065 | 1.77E-05 | 0.065 |
| 18 | ECG | 5087113 | 4 | 17.7 | 3.74E-03 | 0.035 | 2.92E-04 | 0.048 |
| 19 | EGC | 5112438 | 4 | 27.4 | 1.50E-11 | 0.148 | 1.95E-08 | 0.134 |
| 20 | EGC | 5123257 | 4 | 26.7 | 9.59E-13 | 0.162 | 6.95E-10 | 0.167 |
| 21 | EGC | 5136058 | 4 | 27.6 | 3.43E-11 | 0.143 | 3.60E-09 | 0.146 |
| 22 | EGCG | 5074553 | 4 | 39.5 | 3.26E-14 | 0.210 | 1.86E-12 | 0.221 |
| 23 | EGCG | 5123463 | 4 | 28.9 | 3.38E-17 | 0.250 | 1.09E-13 | 0.266 |
| 24 | EGCG | 5134853 | 4 | 37.6 | 7.03E-14 | 0.205 | 2.00E-09 | 0.170 |
| 25 | EGCG | 5136554 | 4 | 38.7 | 1.46E-16 | 0.242 | 2.08E-11 | 0.222 |
| 26 | EGCG | 100011901\|F\|0 | 4 | 38.7 | 1.24E-04 | 0.066 | 3.13E-03 | 0.039 |
| 27 | TF1 | 5064764 | 4 | 28.8 | 2.35E-10 | 0.149 | 5.78E-06 | 0.088 |
| 28 | TF1 | 5106352 | 4 | 26.0 | 1.95E-04 | 0.061 | 8.45E-03 | 0.032 |
| 29 | TF1 | 5123463 | 4 | 28.9 | 1.33E-10 | 0.153 | 2.66E-06 | 0.097 |
| 30 | TF1 | 5136945 | 4 | 28.1 | 1.69E-11 | 0.166 | 8.20E-07 | 0.107 |
| 31 | TF2 | 5063001 | 4 | 30.3 | 9.76E-04 | 0.050 | 4.22E-03 | 0.035 |
| 32 | TF2 | 5064764 | 4 | 28.8 | 1.85E-08 | 0.122 | 7.36E-05 | 0.063 |
| 33 | TF2 | 5106022 | 4 | 29.1 | 2.99E-09 | 0.134 | 3.06E-04 | 0.052 |
| 34 | TF4 | 5064764 | 4 | 28.8 | 8.64E-06 | 0.082 | 1.33E-03 | 0.042 |
| 35 | TF4 | 5106022 | 4 | 29.1 | 1.38E-06 | 0.094 | 1.57E-03 | 0.040 |
| 36 | TF4 | 5136945 | 4 | 28.1 | 6.26E-07 | 0.099 | 1.59E-04 | 0.060 |
| 37 | CL | 5132432 | 7 | 72.7 | 1.31E-03 | 0.047 | 7.30E-04 | 0.049 |
| 38 | ECG | 5059563 | 12 | 49.0 | 8.91E-09 | 0.111 | 2.29E-05 | 0.068 |
| 39 | ECG | 5136077 | 12 | 42.8 | 1.91E-07 | 0.094 | 6.34E-03 | 0.027 |
| 40 | ECG | 5136790 | 12 | 50.4 | 7.87E-11 | 0.137 | 1.58E-04 | 0.051 |
| 41 | EGC | 5135536 | 12 | 50.4 | 2.25E-04 | 0.053 | 2.82E-06 | 0.086 |
| 42 | EGCG | 5087581 | 12 | 48.3 | 1.51E-04 | 0.065 | 1.75E-03 | 0.040 |
| 43 | EGCG | 5088456 | 12 | 47.9 | 7.70E-04 | 0.053 | 2.46E-04 | 0.060 |
| 44 | EGCG | 5133629 | 12 | 47.7 | 7.17E-04 | 0.054 | 1.95E-03 | 0.041 |
| 45 | EC | 5085963 | 15 | 25.4 | 3.52E-04 | 0.049 | 9.09E-03 | 0.029 |
| 46 | EC | 5132852 | 15 | 24.8 | 1.09E-04 | 0.056 | 2.50E-03 | 0.036 |
| 47 | EC | 5099958_E-25 | 15 | 25.1 | 1.00E-04 | 0.056 | 6.37E-03 | 0.031 |
| 48 | TF3 | 5083854 | 15 | 15.2 | 3.52E-03 | 0.043 | 2.77E-03 | 0.035 |
| 49 | TF3 | 5136247 | 15 | 16.3 | 1.57E-03 | 0.049 | 5.40E-04 | 0.046 |

<sup>c</sup> Putative QTL identified in General Linear Model and Mixed Linear Model methods using TASSEL software v5.2.43
LG- Linkage group
Marker R<sup>2</sup>- percentage phenotypic variation explained by markers in both GLM (Q) and MLM (Q+K) methods
Q- Population structure matrix
K- Kinship matrix
CAFF- Caffeine; CAT- Catechin; EC- Epicatechin; ECG- Epicatechin gallate; EGC- Epigallocatechin gallate; EGCG- Epigallocatechin gallate; TF1- Theaflavin; TF2- Theaflavin-3-gallate; TF3- Theaflavin-3'-gallate; TF4- Theaflavin-3,3'-digallate; CL- Colour; BRT- Brightness; AST- Astringency; BRK- Briskness; AR- Aroma; RWC- percent relative water content

A total of 49 DArT markers were associated with 16 different phenotypic traits in six out of 15 linkage groups (Table 3). Since, all 1,421 DArT markers in the genotypic dataset had a known map location, the loci associated with particular traits could be allocated to a linkage group. The numbers of marker loci associated with traits together with their location and positions in different linkage groups at a test level of p< 0.01 in the two test models are shown in Table 3. Choosing a significance level of p< 0.01 which involves multiple testing corrections in both GLM (Q) and MLM (Q+K) models, 32 DArT marker loci were detected for all individual catechins (C, EC, ECG, EGC, EGCG), scattered over five linkage groups namely LG01, LG02, LG04, LG12 and LG15 (Table 3). For caffeine trait, three DArT markers were only located in LG02 while 11 DArT markers for all individual theaflavins traits (TF1, TF2, TF3, TF4) were detected in LG04 and LG15 (Table 3). The trait for tea liquor colour which is an indicator for black tea quality in terms of tea tasters’ score appeared to be associated with only one DArT marker locus at the significance level of p< 0.01 in LG07 (Table 3). Drought tolerance trait (%RWC) was also associated with only one DArT marker locus detected in LG02 (Table 3). The majority of the DArT markers loci associated with the traits were found in LG04, LG12, LG15, LG02, and LG01. The three DArT markers associated with catechin trait in LG04, namely, 5136945, 5064764 and 5134490 had the highest proportion of the percent explained phenotypic variation of 41%, 38%, and 25%, respectively. The DArT markers that also showed high proportion of percent explained phenotypic variation were also observed in LG04 for qEGCG, qEGC, and qTF1 traits, respectively. In this study, a pleiotropic locus which is associated with a single locus affecting two or more distinct phenotypic traits was observed. Several pleiotropic loci were identified including DArT markers 5136945, 5112438, 5136058, 5123257, 5106022 and 5064764 which were associated with qC, qEC, qEGC, qTF1, qTF2 and qTF4 phenotypic traits (Table 3). All the pleiotropic loci detected and associated with the respective phenotypic traits were located in LG04.

#### A comparison of linkage mapping and association mapping

The putative QTLs detected using IM, and MQM mapping in MapQTL 6.0 were compared with marker-trait association results obtained using GLM (Q) and MLM (Q+K) models in AM. The significant marker-trait association results obtained using AM were similar to with the results obtained using MapQTL 6.0 (Tables 4 and 5). The IM analysis produced 14 significant QTLs (LOD > 3.0) that were also found to be associated with the phenotypic traits in AM analysis (Table 4). From the 14 QTLs, a total of 11 markers were associated with four individual catechins (qCAT, qEC, qECG, and qEGC) and one marker each was associated with qTF2, qCL, and qRWC, respectively. On the other hand, the MQM mapping analysis produced eight significant QTLs (LOD > 3.0) which were also found to be significantly associated with the phenotypic traits in AM analysis (Table 5). Of these, five markers were associated with four individual catechins (qCAT, qEC, qECG, and qEGC) and one marker each was associated with qTF2, qCL, and qRWC, respectively. Furthermore, the four mapping models were compared with each other to find markers that could associate with phenotypic traits in all four mapping models (Table 6). A total of 13 markers were found to be associated with four individual catechins (qCAT, qEC, qECG, and qEGC) and one marker each was associated with qTF2, qCL, and qRWC, respectively. Most of the markers that were associated with different phenotypic traits were only located in LG01, LG02, LG04, LG12 and LG15 with a majority of markers showing both positive and negative male additive effects.

**Table 4.** Association mapping and linkage mapping results for putative QTLs in black tea quality and drought tolerance traits in St.504 and St.524 using GLM (Q), MLM (Q + K) methods (p < 0.01) in TASSEL software v5.2.43 and interval mapping in MapQTL 6.0

| Nr | <sup>ac</sup>Trait | Marker | LG | Position<br>(cM) | p-Value | Marker R<sup>2</sup><br>GLM (Q) | p-Value | Marker R<sup>2</sup><br>MLM (Q+K) |
| --- | --- | --- | --- | --- | --- | --- | --- | --- |
| 1 | ECG | 5128890 | 1 | 96.4 | 1.18E-03 | 0.042 | 5.31E-03 | 0.031 |
| 2 | EGC | 5133866 | 1 | 87.1 | 8.94E-03 | 0.030 | 6.62E-04 | 0.046 |
| 3 | RWC | 5136794 | 2 | 60.9 | 8.58E-03 | 0.032 | 3.02E-03 | 0.033 |
| 4 | CAT | 5063001 | 4 | 30.3 | 2.34E-09 | 0.098 | 6.67E-08 | 0.109 |
| 5 | EC | 5123257 | 4 | 26.7 | 3.89E-08 | 0.101 | 7.16E-07 | 0.089 |
| 6 | EC | 5134490 | 4 | 26.4 | 5.15E-19 | 0.228 | 6.21E-15 | 0.246 |
| 7 | ECG | 5087113 | 4 | 17.7 | 3.74E-03 | 0.035 | 2.92E-04 | 0.048 |
| 8 | EGC | 5123257 | 4 | 26.7 | 9.59E-13 | 0.162 | 6.95E-10 | 0.167 |
| 9 | TF2 | 5063001 | 4 | 30.3 | 9.76E-04 | 0.050 | 4.22E-03 | 0.035 |
| 10 | CL | 5132432 | 7 | 72.7 | 1.31E-03 | 0.047 | 7.30E-04 | 0.049 |
| 11 | ECG | 5136790 | 12 | 50.4 | 7.87E-11 | 0.137 | 1.58E-04 | 0.051 |
| 12 | EGC | 5135536 | 12 | 50.4 | 2.25E-04 | 0.053 | 2.82E-06 | 0.086 |
| 13 | EC | 5085963 | 15 | 25.4 | 3.52E-04 | 0.049 | 9.09E-03 | 0.029 |
| 14 | EC | 5099958_E-25 | 15 | 25.1 | 1.00E-04 | 0.056 | 6.37E-03 | 0.031 |

<sup>ac</sup> Putative QTL identified in both Interval Mapping (MapQTL 6.0), General Linear Model and Mixed Linear Model methods (TASSEL software v5.2.43)
Marker R<sup>2</sup>- percentage phenotypic variation explained by markers in GLM and MLM
CAT- Catechin; EC- Epicatechin; ECG- Epicatechin gallate; EGC- Epigallocatechin gallate; TF2- Theaflavin-3-gallate; CL- Colour; RWC- Percent relative water content

**Table 5.** Association mapping and linkage mapping results for putative QTLs in black tea quality and drought tolerance traits in St.504 and St.524 using GLM (Q), MLM (Q + K) methods (p < 0.01) in TASSEL software v5.2.43 and multiple QTL model mapping in MapQTL 6.0

| Nr | <sup>bc</sup>Trait | Marker | LG | Position<br>(cM) | p-Value | Marker R<sup>2</sup><br>GLM (Q) | p-Value | Marker R<sup>2</sup><br>MLM (Q+K) |
| --- | --- | --- | --- | --- | --- | --- | --- | --- |
| 1 | ECG | 5128890 | 1 | 96.4 | 1.18E-03 | 0.042 | 5.31E-03 | 0.031 |
| 2 | EGC | 5133866 | 1 | 87.1 | 8.94E-03 | 0.030 | 6.62E-04 | 0.046 |
| 3 | RWC | 5136794 | 2 | 60.9 | 8.58E-03 | 0.032 | 3.02E-03 | 0.033 |
| 4 | CAT | 5063001 | 4 | 30.3 | 2.34E-09 | 0.098 | 6.67E-08 | 0.109 |
| 5 | TF2 | 5063001 | 4 | 30.3 | 9.76E-04 | 0.050 | 4.22E-03 | 0.035 |
| 6 | CL | 5132432 | 7 | 72.7 | 1.31E-03 | 0.047 | 7.30E-04 | 0.049 |
| 7 | ECG | 5136790 | 12 | 50.4 | 7.87E-11 | 0.137 | 1.58E-04 | 0.051 |
| 8 | EC | 5085963 | 15 | 25.4 | 3.52E-04 | 0.049 | 9.09E-03 | 0.029 |

<sup>bc</sup> Putative QTL identified in both multiple QTL model mapping (MapQTL 6.0), General Linear Model and Mixed Linear Model methods (TASSEL software v5.2.43)
Marker R<sup>2</sup>- percentage phenotypic variation explained by markers in GLM and MLM
CAT- Catechin; EC- Epicatechin; ECG- Epicatechin gallate; EGC- Epigallocatechin gallate; TF2- Theaflavin-3-gallate; CL- Colour; RWC- Percent relative water content

**Table 6.** Association mapping and linkage mapping results for putative QTLs in black tea quality and drought tolerance traits in St.504 and St.524 using GLM (Q), MLM (Q + K) methods (p < 0.01) in TASSEL software v5.2.43 and interval mapping and multiple QTL model mapping in MapQTL 6.0

| Nr | <sup>abc</sup> Trait | Marker | LG | Position (cM) | p-Value | Marker R<sup>2</sup> GLM (Q) | p-Value | Marker R<sup>2</sup> MLM (Q+K) | LOD | PEV | Af | Am | D | Allelic effects |
| --- | --- | --- | --- | --- | --- | --- | --- | --- | --- | --- | --- | --- | --- | --- |
| 1 | ECG | 5128890 | 1 | 96.4 | 1.18E-03 | 0.042 | 5.31E-03 | 0.031 | 6.8 | 11.7 | 0.326 | 0.046 | 0.011 | Af |
| 2 | EGC | 5133866 | 1 | 87.1 | 8.94E-03 | 0.030 | 6.62E-04 | 0.046 | 3.3 | 5.8 | 0.109 | 0.218 | 0.054 | Am |
| 3 | RWC | 5136794 | 2 | 60.9 | 8.58E-03 | 0.032 | 3.02E-03 | 0.033 | 4.3 | 7.3 | 0.482 | 1.13 | 0.283 | Am |
| 4 | CAT | 5063001 | 4 | 30.3 | 2.34E-09 | 0.098 | 6.67E-08 | 0.109 | 5.3 | 4.1 | -17.6 | -53.45 | -13.37 | Am |
| 5 | EC | 5123257 | 4 | 26.7 | 3.89E-08 | 0.101 | 7.16E-07 | 0.089 | 13.4 | 21.1 | 1.703 | 43.81 | 10.95 | Am |
| 6 | ECG | 5087113 | 4 | 17.7 | 3.74E-03 | 0.035 | 2.92E-04 | 0.048 | 3.8 | 5.8 | -2891.86 | 3715.23 | 928.81 | Af; Am |
| 7 | EGC | 5123257 | 4 | 26.7 | 9.59E-13 | 0.162 | 6.95E-10 | 0.167 | 13.4 | 21.1 | 1.703 | 43.81 | 10.95 | Am |
| 8 | TF2 | 5063001 | 4 | 30.3 | 9.76E-04 | 0.050 | 4.22E-03 | 0.035 | 5.3 | 4.1 | -17.604 | -53.47 | -13.37 | Am |
| 9 | CL | 5132432 | 7 | 72.7 | 1.31E-03 | 0.047 | 7.30E-04 | 0.049 | 3.2 | 6.5 | -0.138 | -0.118 | -0.029 | Af; Am |
| 10 | ECG | 5136790 | 12 | 50.4 | 7.87E-11 | 0.137 | 1.58E-04 | 0.051 | 15 | 23.3 | -22.37 | 60.82 | 15.21 | Am |
| 11 | EGC | 5135536 | 12 | 50.4 | 2.25E-04 | 0.053 | 2.82E-06 | 0.086 | 15 | 23.2 | -22.32 | 60.81 | 15.20 | Am |
| 12 | EC | 5085963 | 15 | 25.4 | 3.52E-04 | 0.049 | 9.09E-03 | 0.029 | 4.4 | 7.5 | 11.67 | -24.47 | -6.12 | Am |
| 13 | EC | 5099958_E-25 | 15 | 25.1 | 1.00E-04 | 0.056 | 6.37E-03 | 0.031 | 4.4 | 7.4 | 11.69 | -24.1 | -6.03 | Am |

<sup>abc</sup> Putative QTL identified using both Interval Mapping and Multiple QTL Model Mapping (MapQTL 6.0) and General Linear Model and Mixed Linear Model method (TASSEL software v5.2.43)
Marker R<sup>2</sup>- percentage phenotypic variation explained by markers in GLM and MLM
LG- Linkage group
LOD- Logarithm of odds ratio for putative QTL
PEV- percentage phenotypic variation explained by markers in IM and MQM
Af- Female additivity
Am- Male additivity
D- Dominance
CAT- Catechin; EC- Epicatechin; ECG- Epicatechin gallate; EGC- Epigallocatechin gallate; TF2- Theaflavin-3-gallate; CL- Colour; RWC- Percent relative water content

#### Functional annotation of putative QTLs in linkage and association mapping

The putative QTLs detected in both IM, and MQM mapping were identified by the BLAST algorithm, searched on the reference tea genome (NCBI) and assigned functions on Blast2GO database. A total of 53 QTLs were detected in both interval and MQM mapping methods, of which 45 proteins were assigned functions. An additional six markers in LG02, LG04, and LG12 were found to be associated with caffeine, EC, ECG and EGC traits in AM (Table 7). The six additional QTLs detected in AM were also functionally annotated using the above databases. The putative candidate genes identified were involved in purine or thiamine biosynthesis, phenylalanine biosynthesis, tyrosine and tryptophan biosynthesis, carbon fixation (photosynthesis) and abiotic stress. The putative QTL, qCaffeine in LG02 was putatively annotated N-(5’phosphoribosyl) anthranilate (PRA) isomerase enzyme. The two other putative QTLs identified for qEC and qEGC located in LG04 were annotated as phosphoribulokinase or uridine kinase family proteins. Two putative QTLs, for qEC in LG04, were annotated as autophagy-related protein 11 and protein tyrosine kinase, respectively.

**Table 7.**
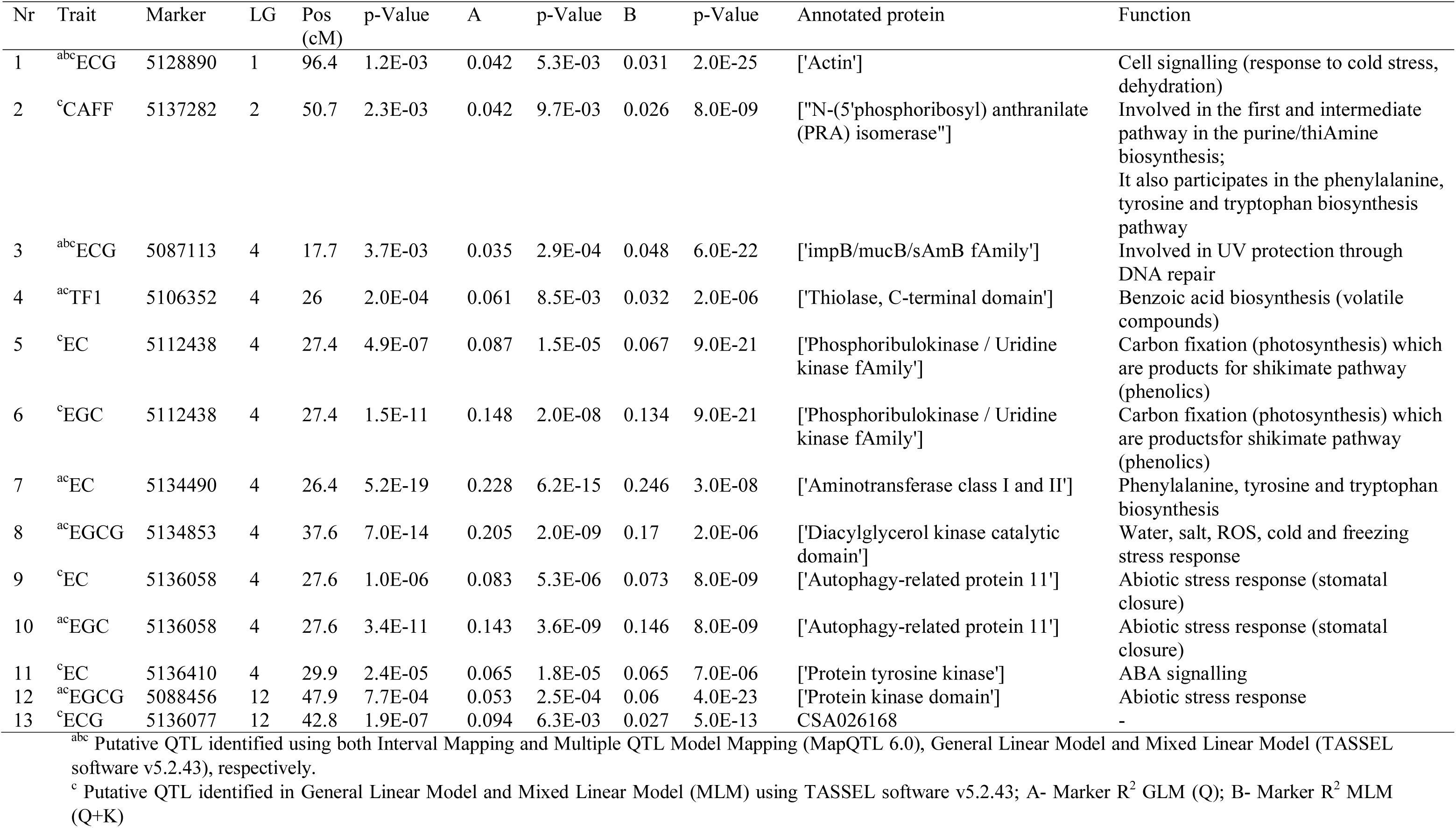
Association mapping, linkage mapping and functional annotation protein of putative QTLs in black tea quality traits in St.504 and St.524 based on IM and MLM (Q + K) method (p < 0.01) in MapQTL 6.0 and TASSEL software v5.2.43, respectively.

| Nr | Trait | Marker | LG | Pos<br>(cM) | p-Value | A | p-Value | B | p-Value | Annotated protein | Function |
| --- | --- | --- | --- | --- | --- | --- | --- | --- | --- | --- | --- |
| 1 | <sup>abc</sup>ECG | 5128890 | 1 | 96.4 | 1.2E-03 | 0.042 | 5.3E-03 | 0.031 | 2.0E-25 | ['Actin'] | Cell signalling (response to cold stress, dehydration) |
| 2 | <sup>c</sup>CAFF | 5137282 | 2 | 50.7 | 2.3E-03 | 0.042 | 9.7E-03 | 0.026 | 8.0E-09 | ['N-(5'phosphoribosyl) anthranilate (PRA) isomerase"] | Involved in the first and intermediate pathway in the purine/thiAmine biosynthesis;<br>It also participates in the phenylalanine, tyrosine and tryptophan biosynthesis pathway |
| 3 | <sup>abc</sup>ECG | 5087113 | 4 | 17.7 | 3.7E-03 | 0.035 | 2.9E-04 | 0.048 | 6.0E-22 | ['impB/mucB/sAmB fAmily'] | Involved in UV protection through DNA repair |
| 4 | <sup>ac</sup>TF1 | 5106352 | 4 | 26 | 2.0E-04 | 0.061 | 8.5E-03 | 0.032 | 2.0E-06 | ['Thiolase, C-terminal domain'] | Benzoic acid biosynthesis (volatile compounds) |
| 5 | <sup>c</sup>EC | 5112438 | 4 | 27.4 | 4.9E-07 | 0.087 | 1.5E-05 | 0.067 | 9.0E-21 | ['Phosphoribulokinase / Uridine kinase fAmily'] | Carbon fixation (photosynthesis) which are products for shikimate pathway (phenolics) |
| 6 | <sup>c</sup>EGC | 5112438 | 4 | 27.4 | 1.5E-11 | 0.148 | 2.0E-08 | 0.134 | 9.0E-21 | ['Phosphoribulokinase / Uridine kinase fAmily'] | Carbon fixation (photosynthesis) which are productsfor shikimate pathway (phenolics) |
| 7 | <sup>ac</sup>EC | 5134490 | 4 | 26.4 | 5.2E-19 | 0.228 | 6.2E-15 | 0.246 | 3.0E-08 | ['Aminotransferase class I and II'] | Phenylalanine, tyrosine and tryptophan biosynthesis |
| 8 | <sup>ac</sup>EGCG | 5134853 | 4 | 37.6 | 7.0E-14 | 0.205 | 2.0E-09 | 0.17 | 2.0E-06 | ['Diacylglycerol kinase catalytic domain'] | Water, salt, ROS, cold and freezing stress response |
| 9 | <sup>c</sup>EC | 5136058 | 4 | 27.6 | 1.0E-06 | 0.083 | 5.3E-06 | 0.073 | 8.0E-09 | ['Autophagy-related protein 11'] | Abiotic stress response (stomatal closure) |
| 10 | <sup>ac</sup>EGC | 5136058 | 4 | 27.6 | 3.4E-11 | 0.143 | 3.6E-09 | 0.146 | 8.0E-09 | ['Autophagy-related protein 11'] | Abiotic stress response (stomatal closure) |
| 11 | <sup>c</sup>EC | 5136410 | 4 | 29.9 | 2.4E-05 | 0.065 | 1.8E-05 | 0.065 | 7.0E-06 | ['Protein tyrosine kinase'] | ABA signalling |
| 12 | <sup>ac</sup>EGCG | 5088456 | 12 | 47.9 | 7.7E-04 | 0.053 | 2.5E-04 | 0.06 | 4.0E-23 | ['Protein kinase domain'] | Abiotic stress response |
| 13 | <sup>c</sup>ECG | 5136077 | 12 | 42.8 | 1.9E-07 | 0.094 | 6.3E-03 | 0.027 | 5.0E-13 | CSA026168 | - |

<sup>abc</sup> Putative QTL identified using both Interval Mapping and Multiple QTL Model Mapping (MapQTL 6.0), General Linear Model and Mixed Linear Model (TASSEL software v5.2.43), respectively.
<sup>c</sup> Putative QTL identified in General Linear Model and Mixed Linear Model (MLM) using TASSEL software v5.2.43; A- Marker $R^2$ GLM (Q); B- Marker $R^2$ MLM (Q+K)

## Discussion

In this current study, the integration of linkage mapping and AM was used to provide a mutual and precise authentication of QTL, which will enable more reliable results to be obtained. Linkage mapping method has a relatively low genome resolution while AM method is affected by population structure and individual relationships. The integration of linkage mapping and AM has successfully been applied to many plant studies (Fan and Xiong 2003; Jung et al. 2005; Lu et al. 2010; Lou et al. 2015; Cao et al. 2017).

The population structure analysis in discovery mapping panel used in the current study identified two groups or clusters. The dendrogram demonstrated two subspecies of the two parental clones, TRFK 303/577 and GW Ejulu, which are of *C. sinensis* var. *assamica* and *C. sinensis* var. *sinensis* variety, respectively. The neighbour-joining tree for the 261 tea accessions grouped them into two major clusters on the base of Nei’s genetic distance. The phylogenetic relationships for the parental cultivars and their progenies clustering into their respective groups in separate clades based on their parental genotypes were consistent with their genetic backgrounds (Kamunya et al. 2010). The parental clone TRFK 303/577, which is an open-pollinated progeny of clone TRFK 6/8, is high-yielding, drought-tolerant, medium in black tea quality, caffeine, and individual catechins. On the other hand, parental clone GW Ejulu is a low-yielding, high black tea quality and moderate levels of caffeine, but high in total catechins and individual catechin contents (TRFK 2012). The bulk of tea cultivars used for green and black tea production have been derived through individual selection, hybridisation and molecular breeding of *C. sinensis* var. *assamica* and *C. sinensis* var. *sinensi*s varieties to produce tea with desirable characteristics (Yao et al. 2008). Therefore, developing a few markers that have the capability of discriminating the two subspecies and their cultivars is of paramount interest. The results on the linkage mapping for all the phenotypic traits and the QTL positions in different linkage groups in this study were similar as reported in (Koech et al. 2018).

In this study, AM was conducted with three different models, Q, K, and Q+K. The observed –log10 (P) values for QTL deviated from the expected –log10 (P) values in the Q method (GLM), indicated that there might be a few false-positive. However, the addition of genetic relatedness K (kinship) used in a mixed linear model has proven to be more powerful and reduces the number of false-positive associations (Finno et al. 2014). This is because a lot more QTLs were identified in this current study using Q-model as compared with K and Q + K-model. Therefore, there could be higher chances of false-positive or negative errors in AM due to complex population structure (Müller et al. 2011; George 2013). In this study, a mixed linear model approach using the K-matrix or a combination (Q+K) performed better than a GLM (Q). However, the K or the Q + K methods did show to be too strict and resulted in the missing of some possibly useful QTLs. Nearly all the significant QTLs identified in AM using the Q, and Q+K models were in line with those identified in MapQTL 6.0 (Koech et al. 2018). The percentages of phenotypic variation explained (R^2^) by various markers analysed using AM were significant as those analysed using MapQTL 6.0. This is in agreement with previous studies reported in plants on efficiency and robustness of combining linkage mapping and AM for precise identification of QTL (Mammadov et al. 2015; Li et al. 2016).

The six additional QTLs identified in AM were found to be associated with purine or thiamine biosynthesis, phenylalanine, tyrosine and tryptophan biosynthesis, carbon fixation (photosynthesis) and abiotic stress. The putative QTL, qCaffeine located in LG02 was putatively annotated N-(5’phosphoribosyl) anthranilate (PRA) isomerase enzyme. The enzyme is involved in the first and intermediate pathway in the purine or thiamine biosynthesis and tryptophan biosynthesis pathway (Li et al. 1995). Caffeine (1,3,7-trimethylxanthine) and other methylxanthines such as theobromine (3,7-dimethylxanthine) and methyluric acids present in tea are classified as purine alkaloids (Ashihara et al. 2008). Therefore, the putative QTL, qCaffeine could be associated with the biosynthesis of caffeine, theobromine and methyluric acids in tea. Also, phenylalanine, tyrosine, and tryptophan are products of shikimate pathway which is involved in the biosynthesis of plant flavonoids, including catechins (Ghasemzadeh and Ghasemzadeh 2011).

The putative QTLs, qEC, and qEGC in LG04 were annotated as phosphoribulokinase/uridine kinase family proteins which are involved in carbon fixation or photosynthesis in plants (Miziorko 2000). Photosynthesis is an important process in plants for provision of carbon skeletons to the shikimate pathway (Janacek et al. 2009). Catechins are synthesised in the leaves of the tea plant through the acetic-malonic acid and shikimic-cinnamic acid metabolic pathways (Vuong et al. 2011). The chalcone and gallic acid are produced from the shikimic acid pathway, which then produce the different catechins (Chu and Juneja 1997). The two putative QTLs, qEC annotated putatively as autophagy-related protein 11 and protein tyrosine kinase are involved in abiotic stress response through a process of stomatal closure and ABA signalling (Sah et al. 2016; Batoko et al. 2017). Stress conditions can interfere with photosynthetic energy production in plants, which leads to stomatal closure, which inhibits CO_2_ intake and thereby reduces photosynthetic activity in leaves (Chaves et al. 2009). Abscisic acid has been reported to confer abiotic stress-tolerance in crop plants (Sah et al. 2016). In stress conditions like drought, extreme temperature, and high salinity, content in plants increases considerably, inspiring stress-tolerance effects that help plants, adapt, and survive under these stressful situations (Ng et al. 2014). Also, ABA is also required for plant growth and development under non-stress conditions (Nakashima and Yamaguchi-Shinozaki 2013).

## Conclusion

The tea plant is a woody perennial crop with long generation intervals, which usually hinders its genetic improvement by conventional breeding methods. The approach of combining the high QTL detection power of genetic linkage mapping with the high-resolution power of AM allowed identification and precise authentication of putative QTL controlling black tea quality and drought tolerance traits. Based on the two mapping approaches, the putative QTLs associated with caffeine, catechins, theaflavins, tea tasters’ scores, and %RWC detected in linkage mapping and AM were not significantly different except that an additional six more QTLs were detected using AM method. Although, conventional QTL mapping through bi- or multi- parental populations is a powerful method, it suffers from a limited amount of recombination. However, AM can partly overcome the limitation by using a diverse germplasm but may lead to a number of false positive or negative associations. Therefore, the two different methods can be complementary to each other and benefit can be achieved by mitigating the other’s limitations. In this study, the combination of AM and QTL mapping has led to successful identification of QTL with potential application in breeding programs. However, the QTL identified by AM requires further confirmation in bi- or multi-parental populations. This will accelerate tea breeding progress in terms of reduced breeding time, reduced cost, increased efficiency, and precision of selection.

## Author contributions

ZA, SK and RM were involved with the design of the experiment and plant material used. RK was involved in collection of plant material. RK performed the experiments, analyzed samples and interpreted the data. RK wrote the manuscript and revised by RM, SK and ZA. The final manuscript was reviewed and approved by all the authors. The TRI of Kenya and the University of Pretoria had role in study design, data collection and analysis, decision to publish, or preparation of the manuscript.

## Acknowledgement

The authors acknowledge the financial support to conduct this research, and study grants for RK and PM from James Finlay (Kenya) Ltd., George Williamson (Kenya) Ltd., Sotik Tea Company (Kenya) Ltd., Mcleod Russell (Uganda) Ltd., the TRI of Kenya and Southern African Biochemistry and Informatics for Natural Products (SABINA). The *C. sinensis* cultivars used in this study were provided by the TRI of Kenya. Supplementary funding was provided by, the Technology and Human Resources for Industry Programme (THRIP), an initiative of the Department of Trade and Industries of South Africa (dti), the National Research Foundation (NRF) of South Africa, and the University of Pretoria (South Africa).

