## Supplementary Figures 1-4 for "Combined linkage and association mapping of putative QTLs controlling black tea quality and drought tolerance traits"

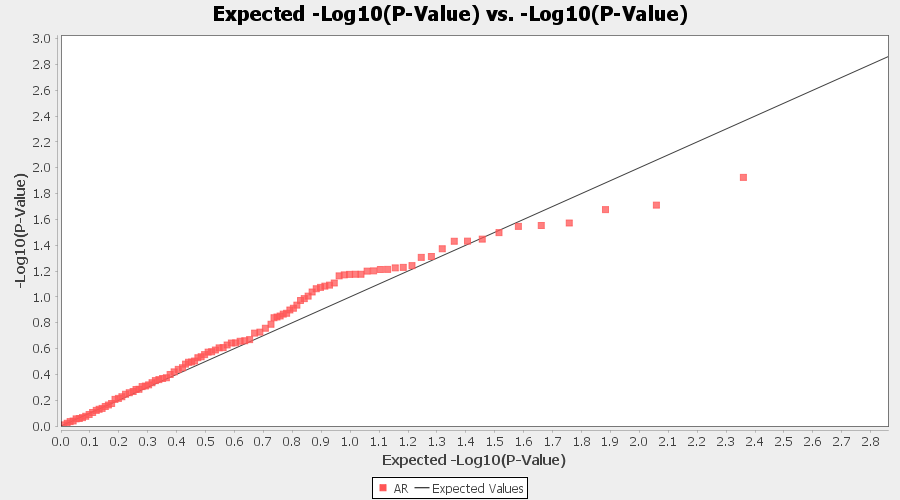

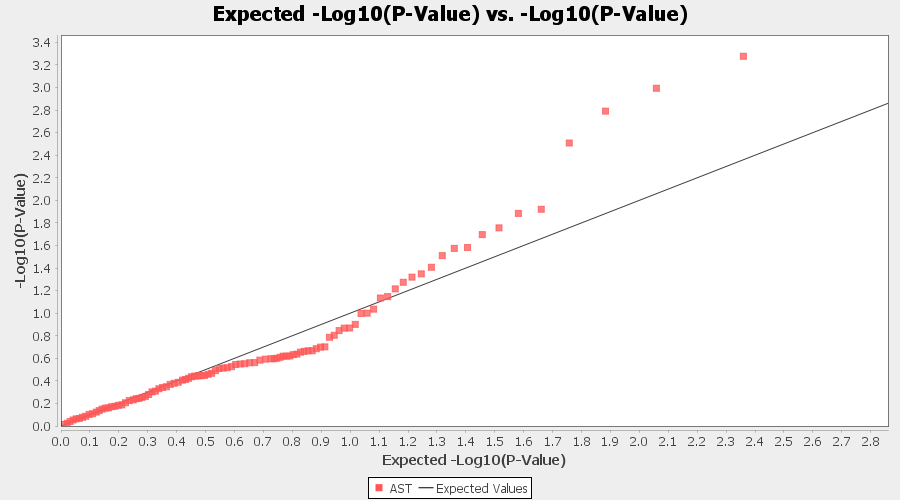

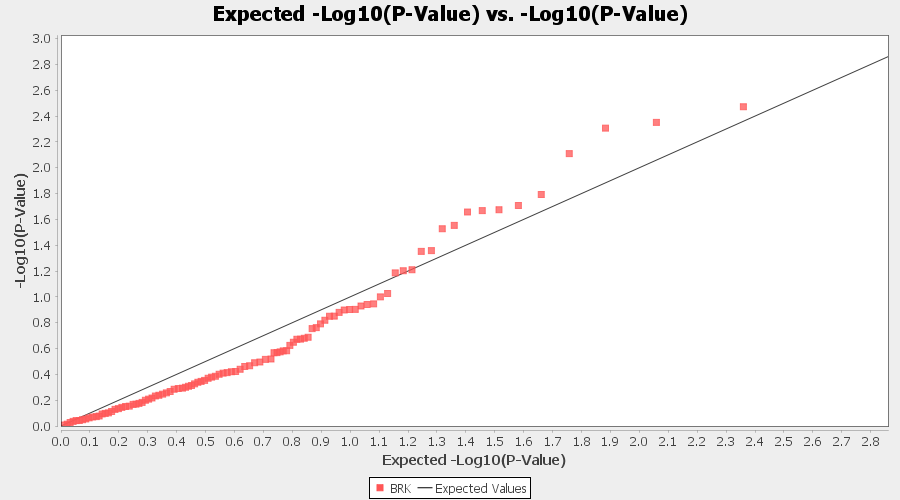

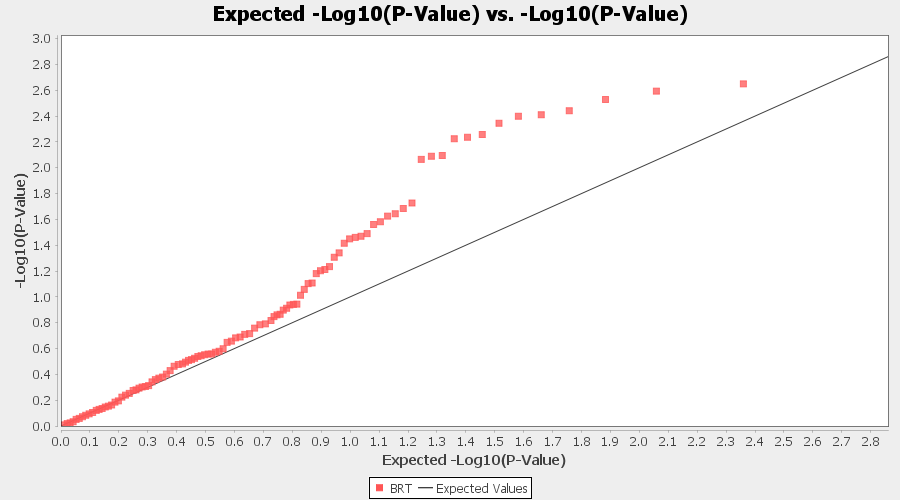

Supplementary Figure 1. Trait-wise association Q-Q plots for GLM. AR- Aroma; AST- Astringency; BRK- Briskness; BRT- Brightness

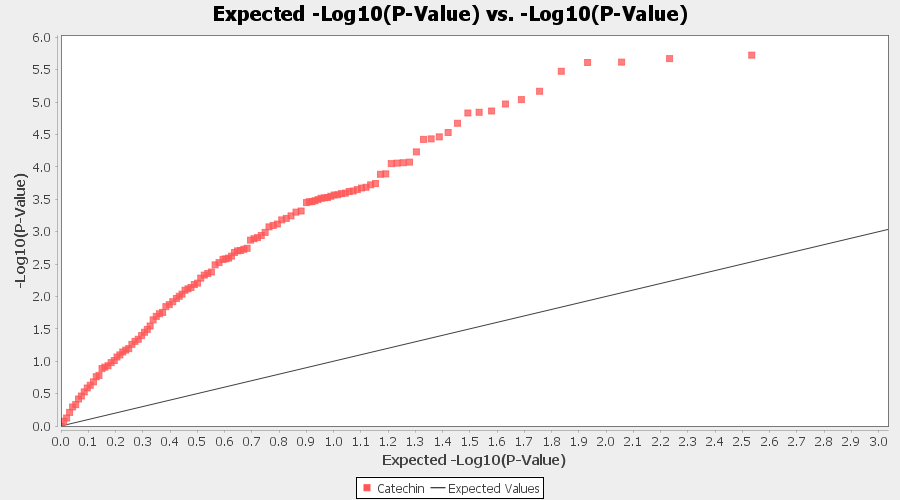

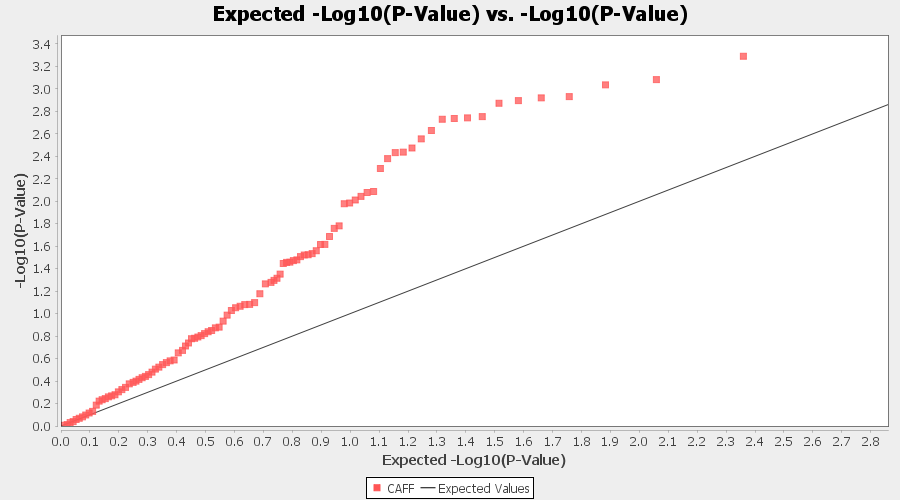

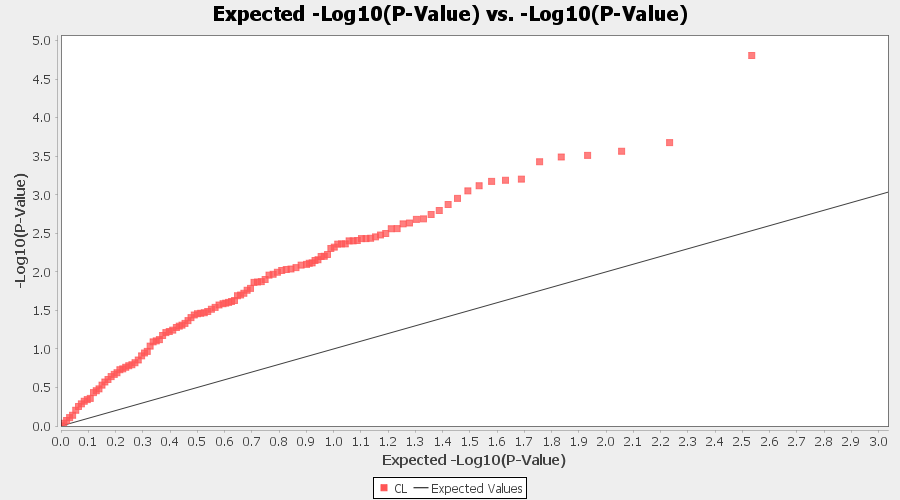

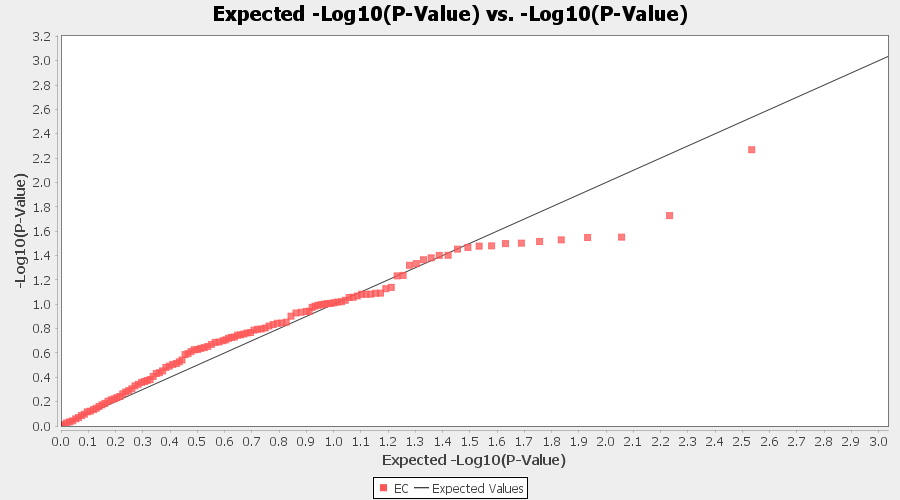

**Continued** Supplementary Figure 1. Trait-wise Q-Q association plots for GLM. C- Catechin; CAFF- Caffeine; CL- Colour; EC- Epicatechin

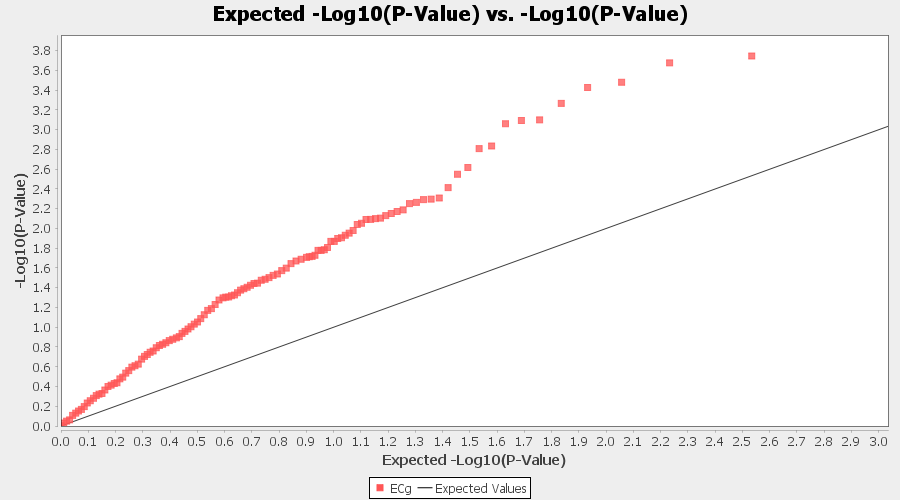

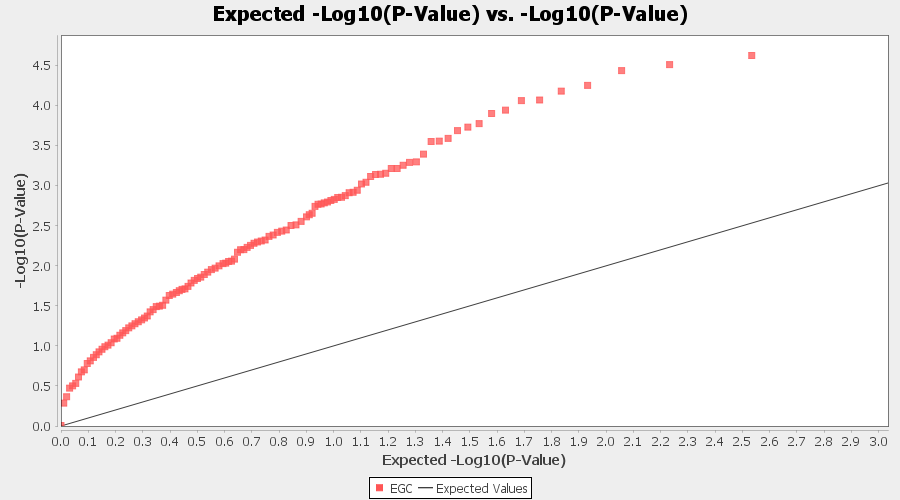

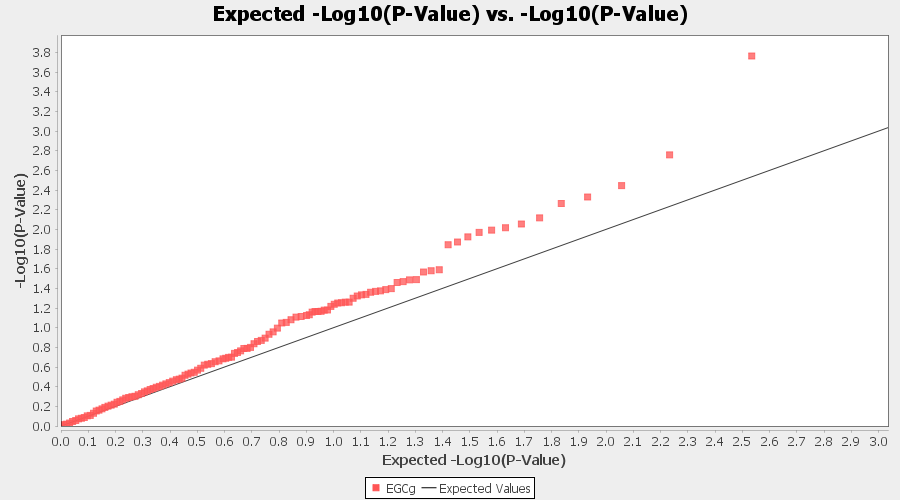

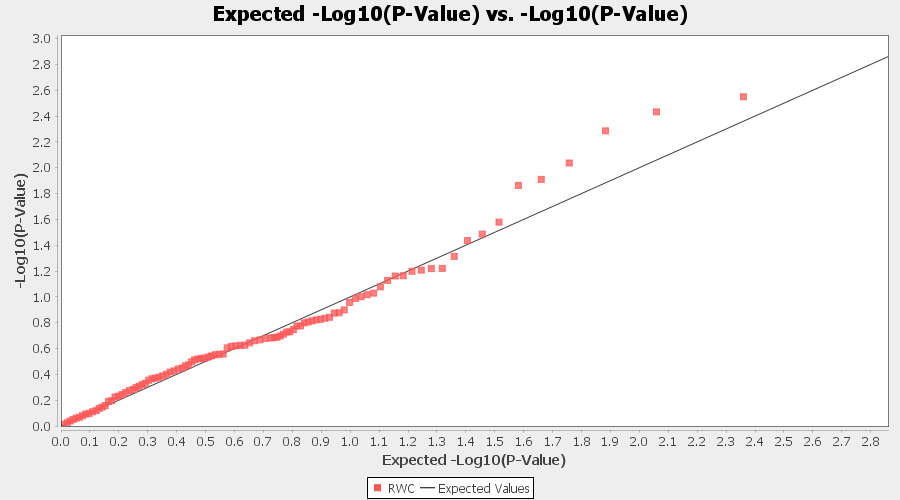

**Continued** Supplementary Figure 1. Trait-wise association Q-Q plots for GLM. ECG- Epicatechin gallate; EGC- Epicatechin gallate; EGCG- Epigallocatechin gallate; RWC- Percent relative water content

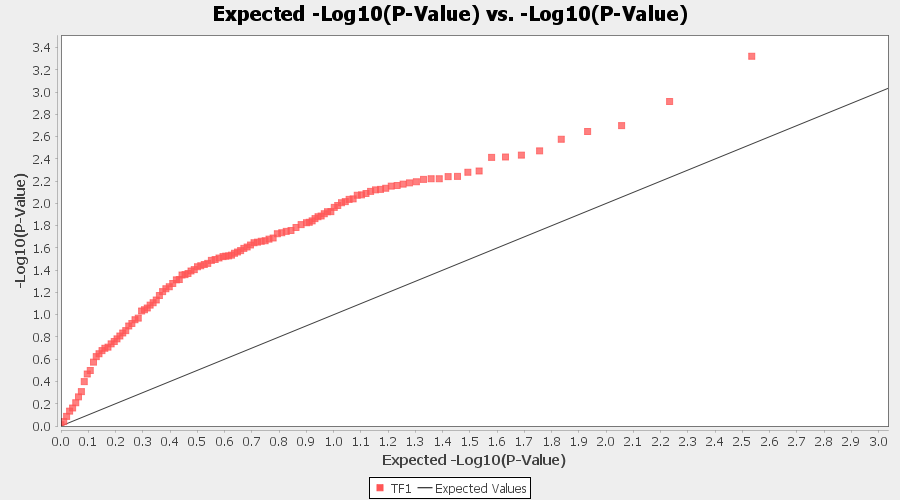

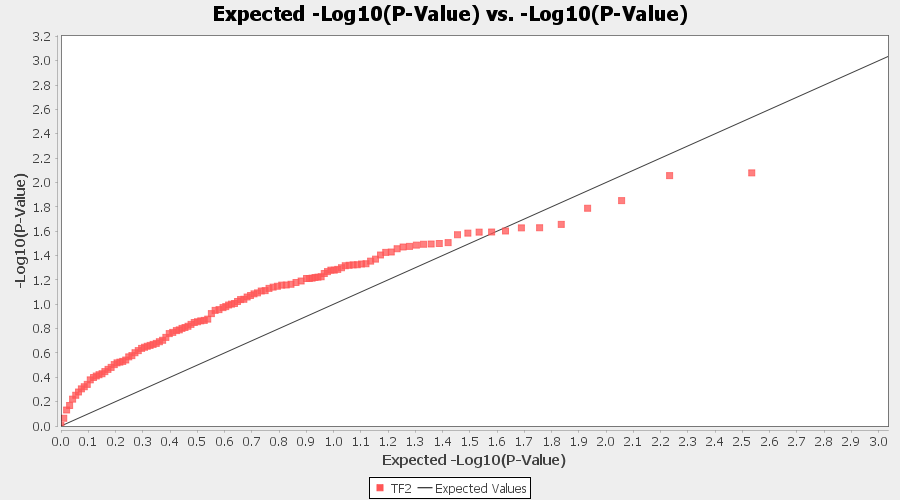

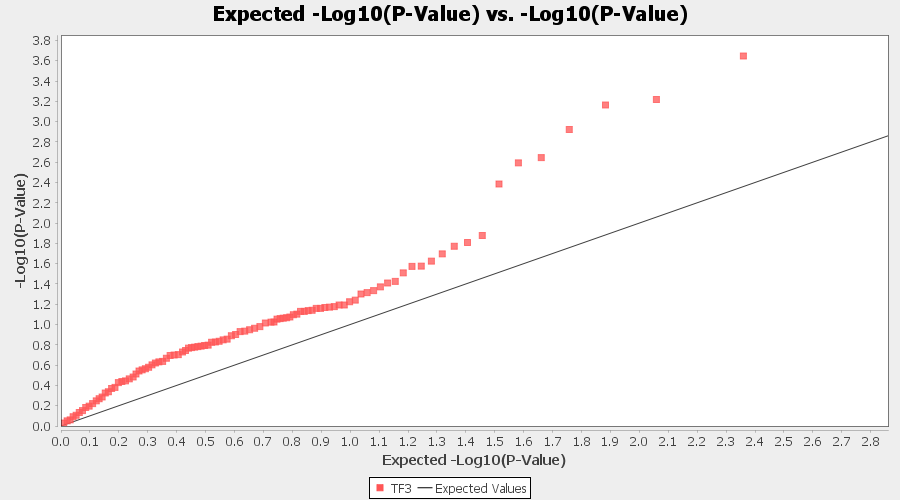

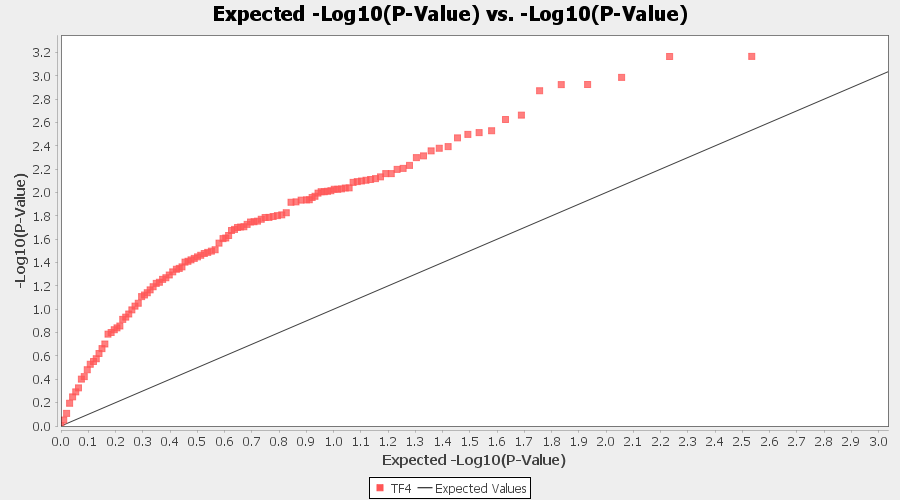

**Continued** Supplementary Figure 1. Trait-wise association Q-Q plots for GLM. TF1- Theaflavin; TF2- Theaflavin-3-gallate; TF3- Theaflavin-3’-gallate; TF4- Theaflavin-3,3’-digallate

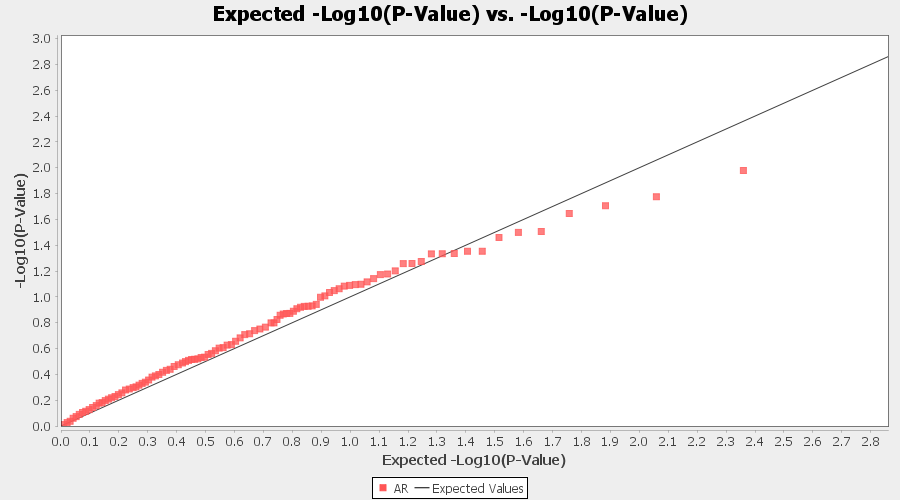

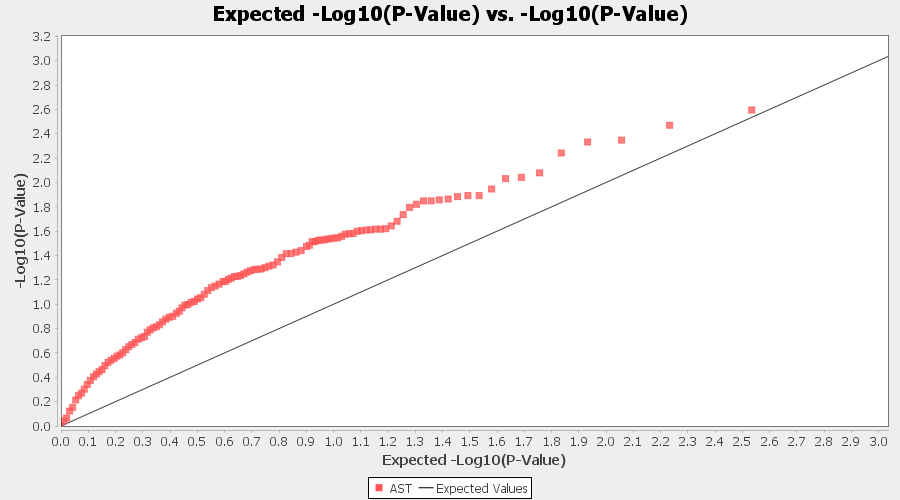

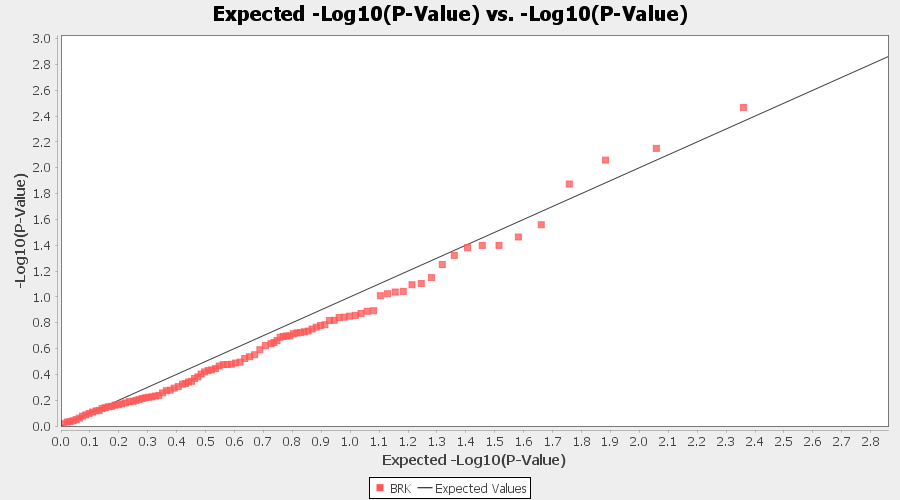

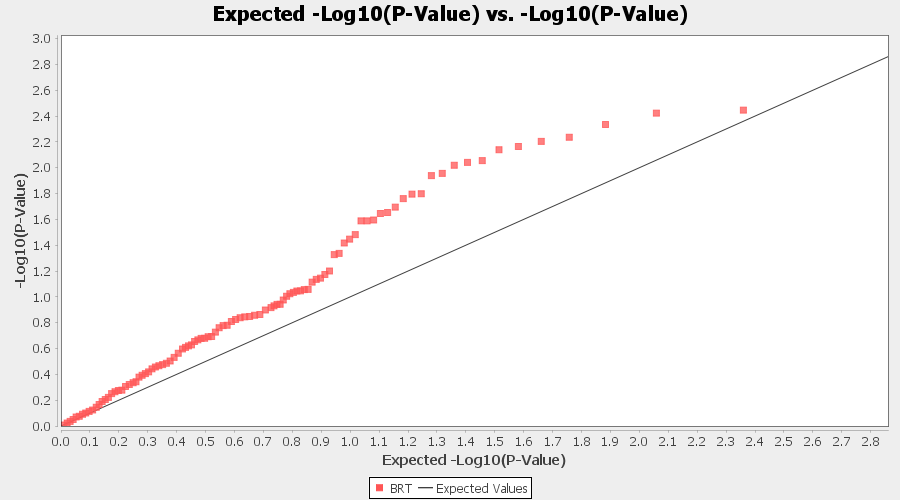

Supplementary Figure 2. Trait-wise association Q-Q plots for GLM (Q). AR- Aroma; AST- Astringency; BRK- Briskness; BRT- Brightness

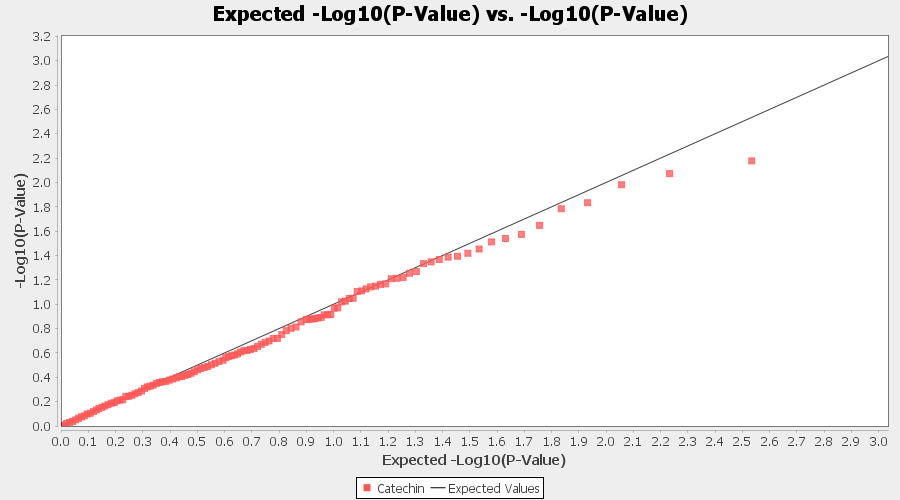

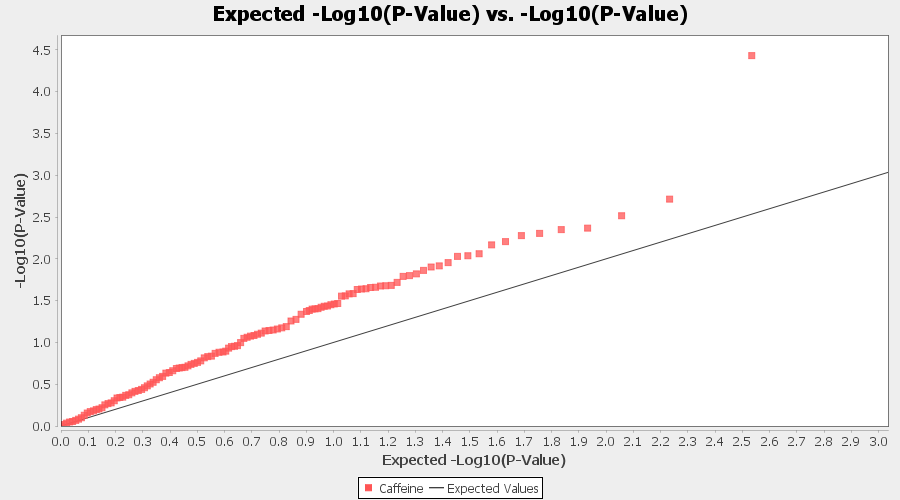

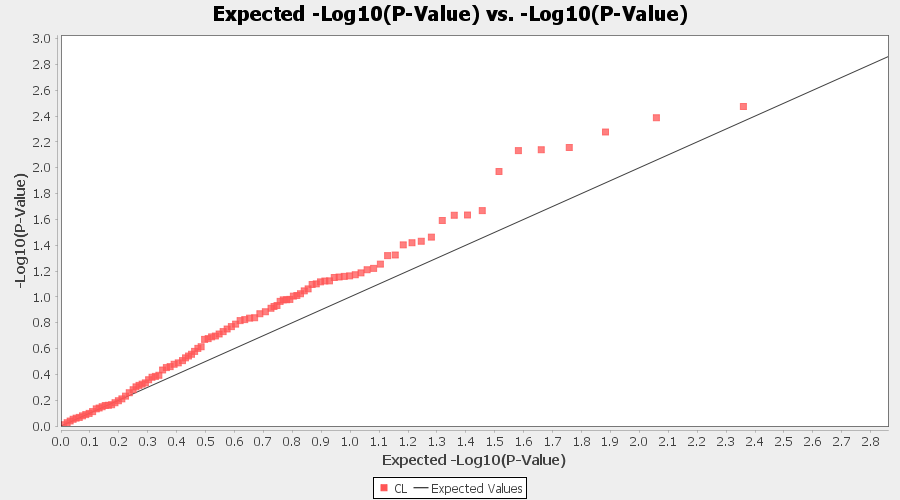

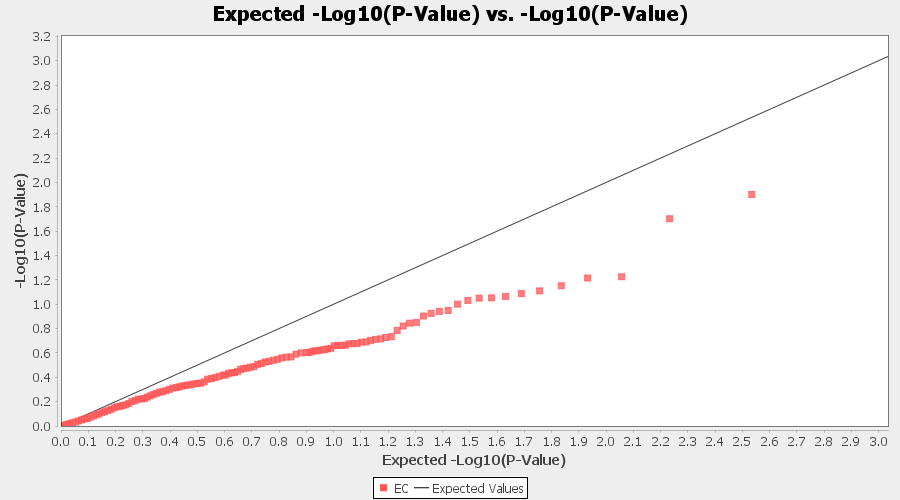

**Continued** Supplementary Figure 2. Trait-wise Q-Q association plots for GLM (Q). C- Catechin; CAFF- Caffeine; CL- Colour; EC- Epicatechin

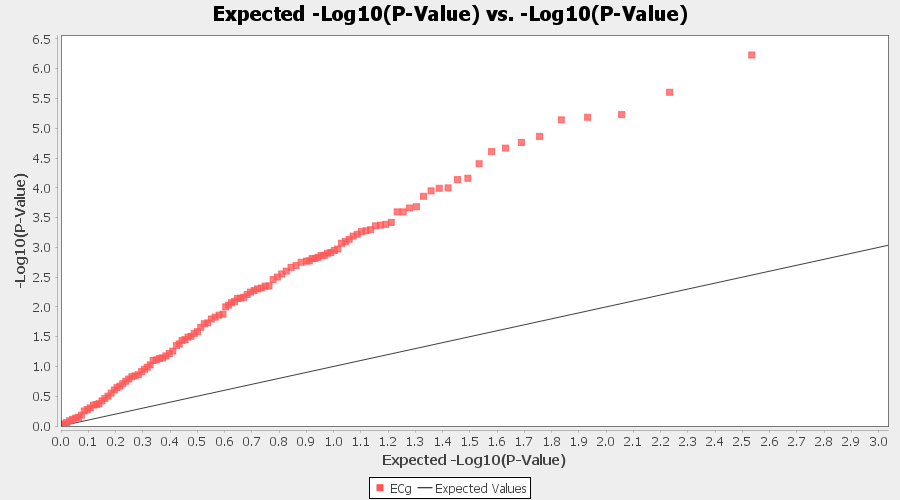

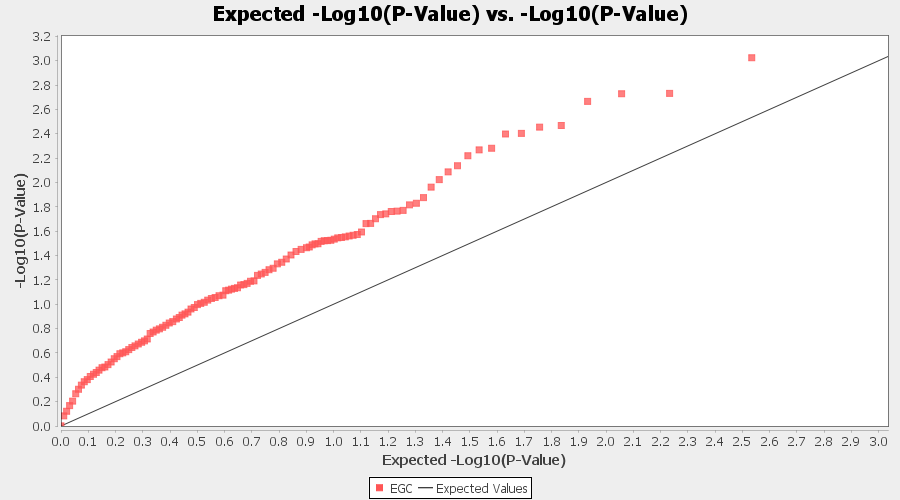

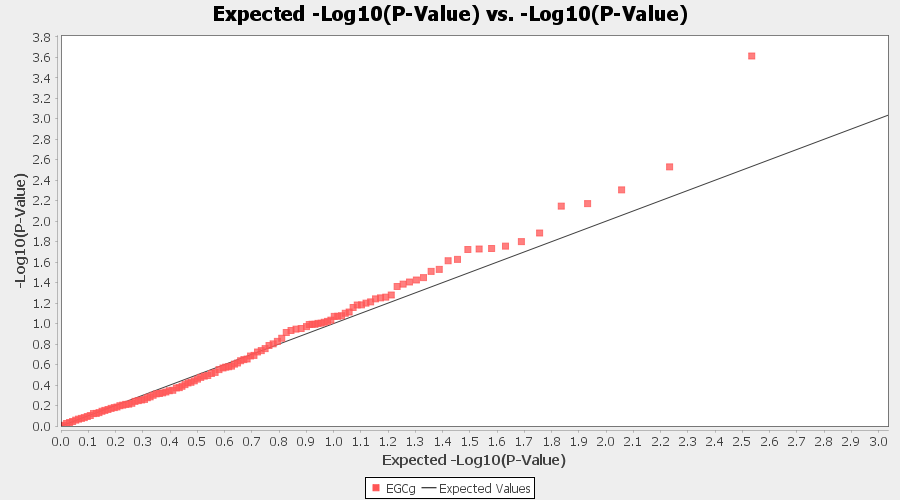

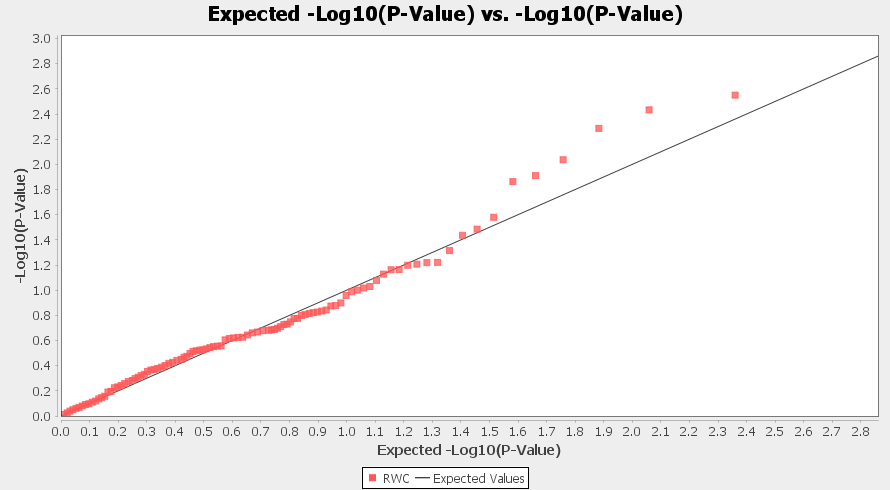

**Continued** Supplementary Figure 2. Trait-wise Q-Q association plots for GLM (Q). ECG- Epicatechin gallate; EGC- Epicatechin gallate; EGCG- Epigallocatechin gallate; RWC- percent relative water content

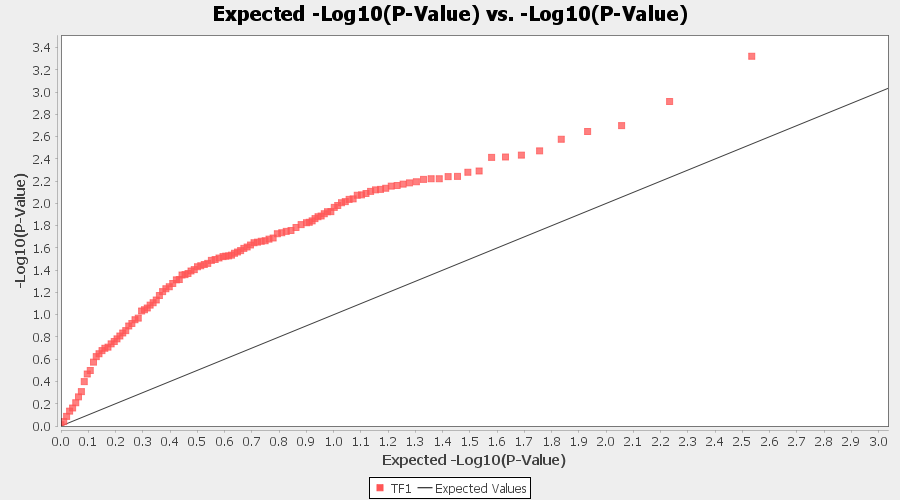

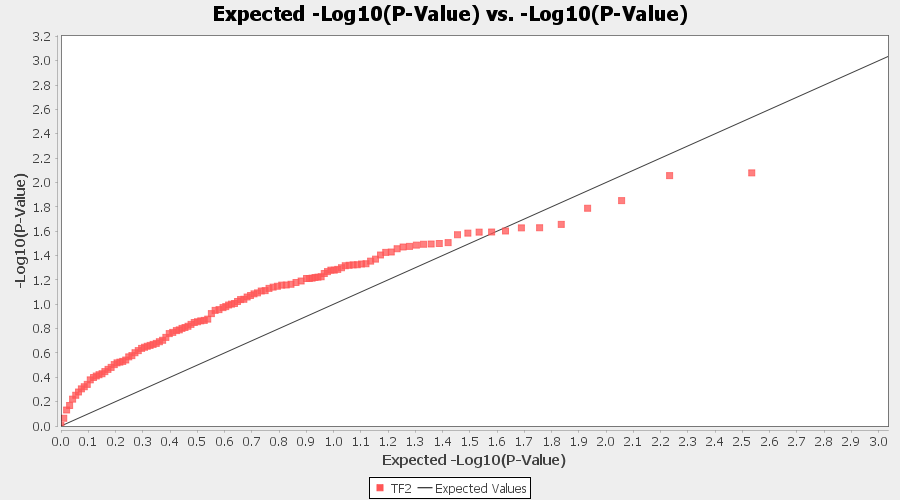

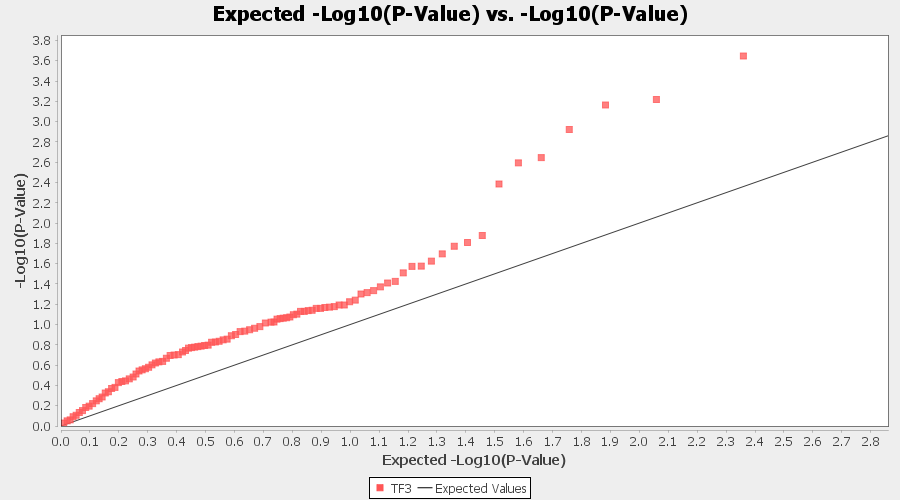

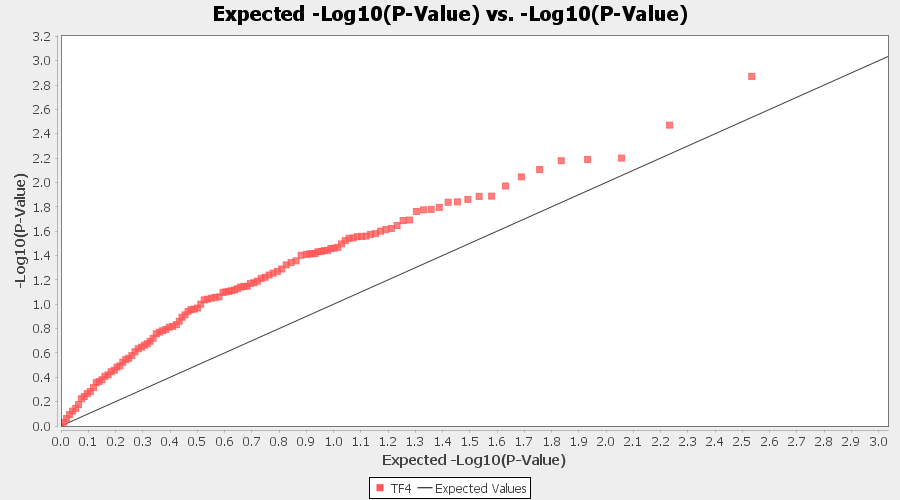

**Continued** Supplementary Figure 2. Trait-wise Q-Q association plots for GLM (Q). TF1- Theaflavin; TF2- Theaflavin-3-gallate; TF3- Theaflavin-3’-gallate; TF4- Theaflavin-3,3’-digallate

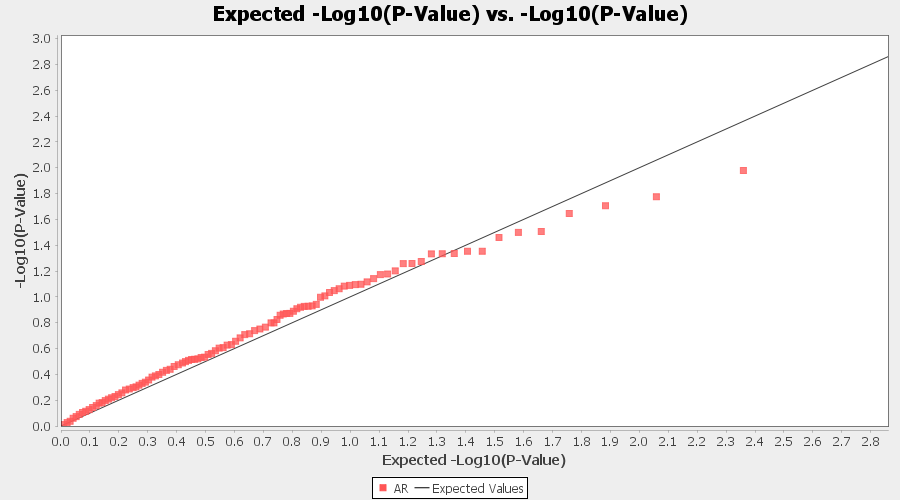

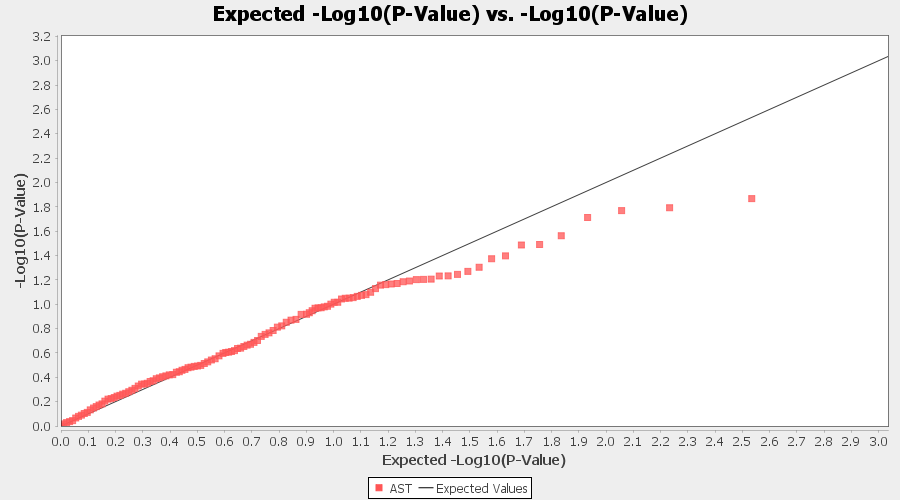

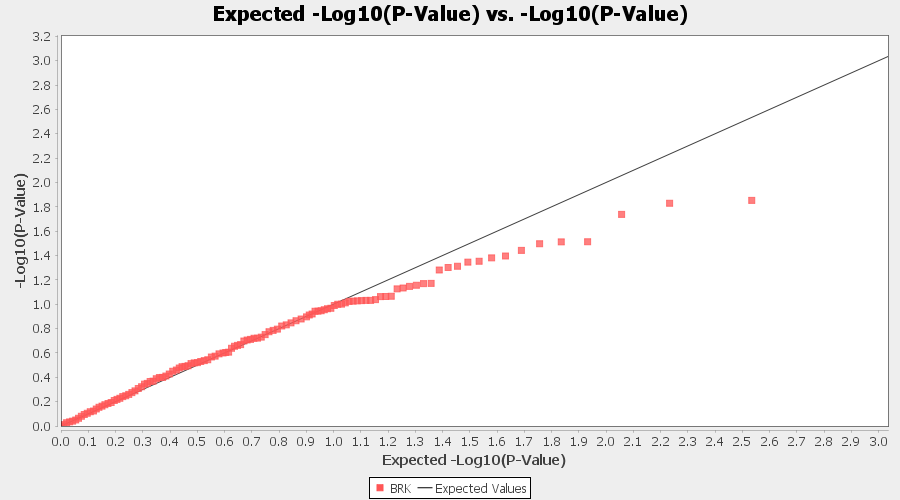

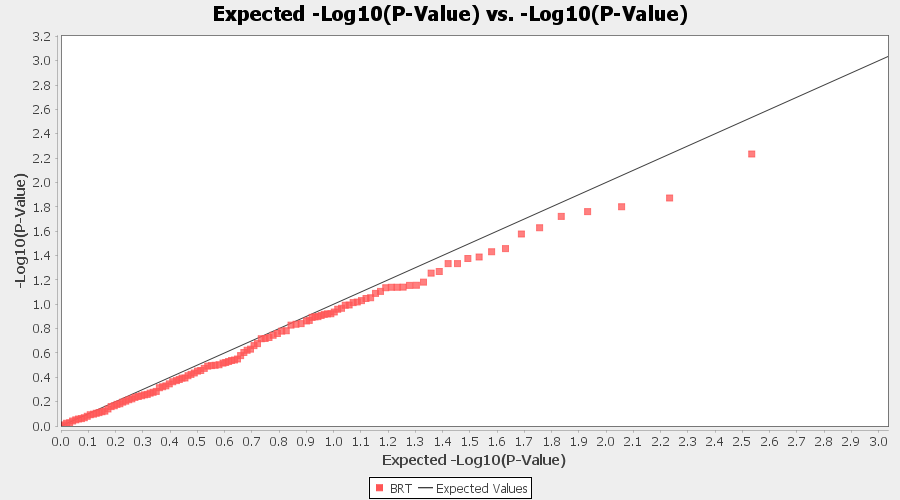

Supplementary Figure 3. Trait-wise Q-Q association plots for mixed linear model MLM (K). AR- Aroma; AST- Astringency; BRK- Briskness; BRT- Brightness

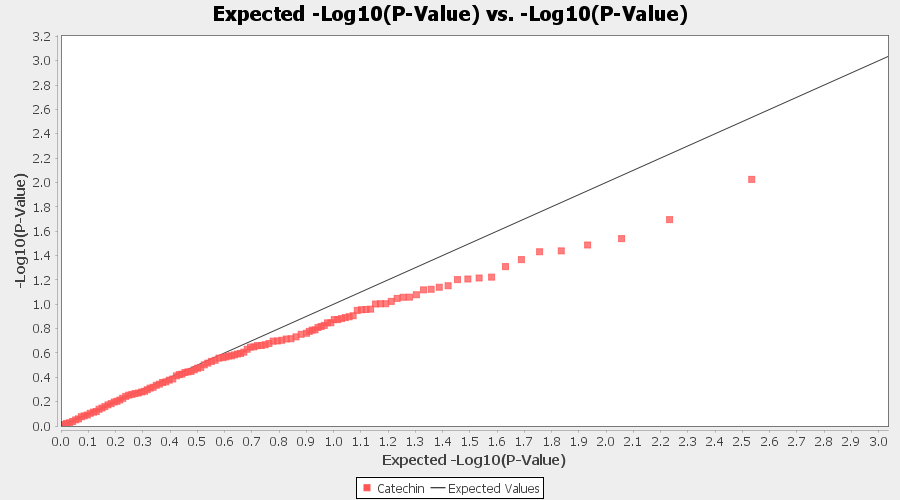

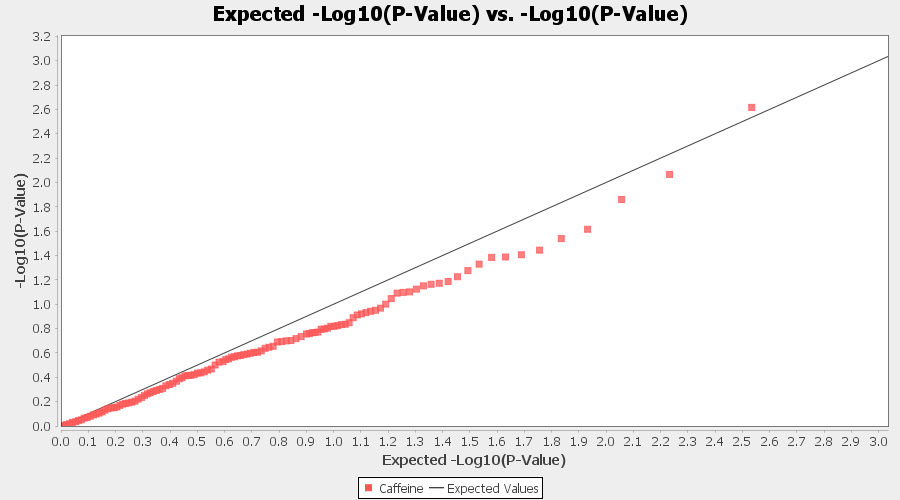

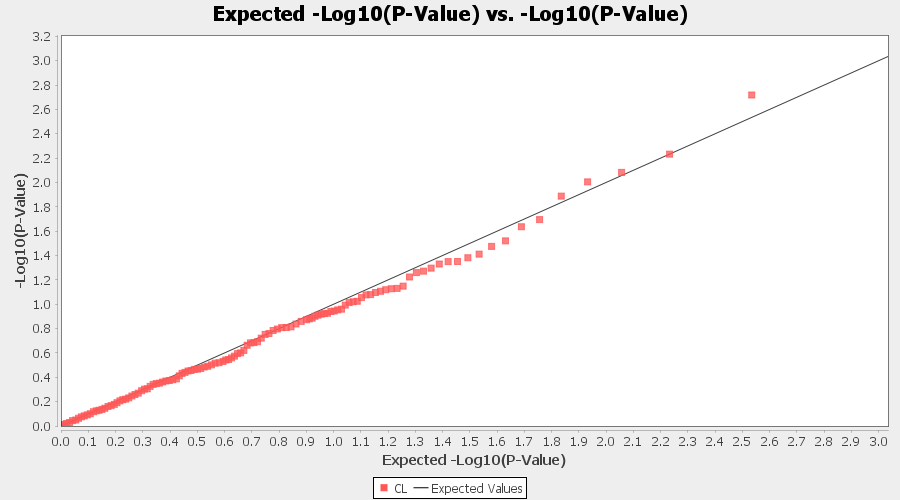

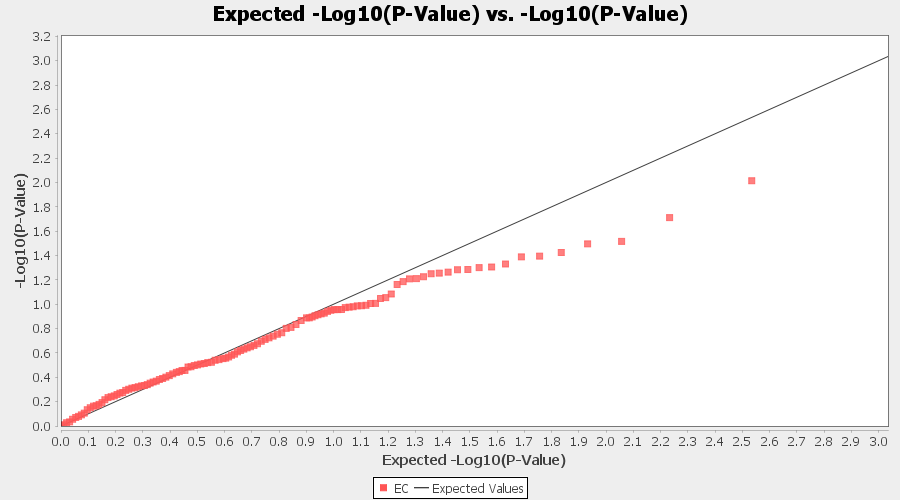

**Continued** Supplementary Figure 3. Trait-wise Q-Q association plots for mixed linear model MLM (K). C- Catechin; CAFF- Caffeine; CL- Colour; EC- Epicatechin

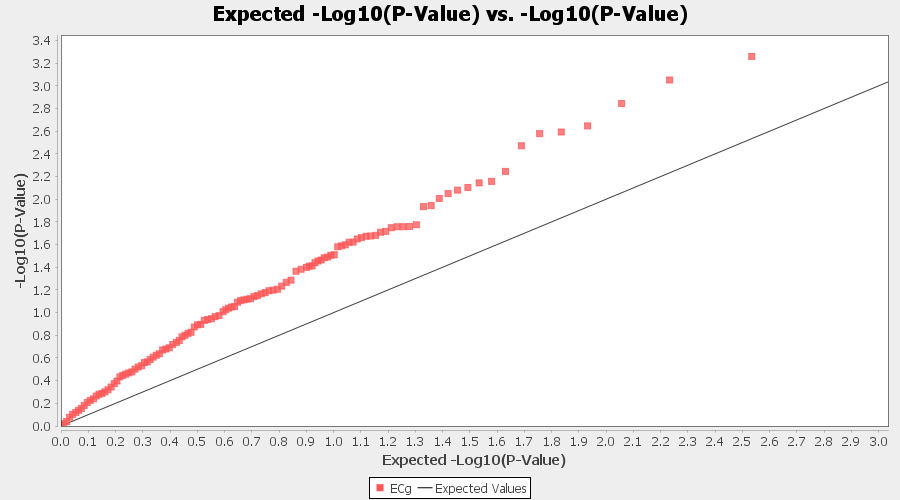

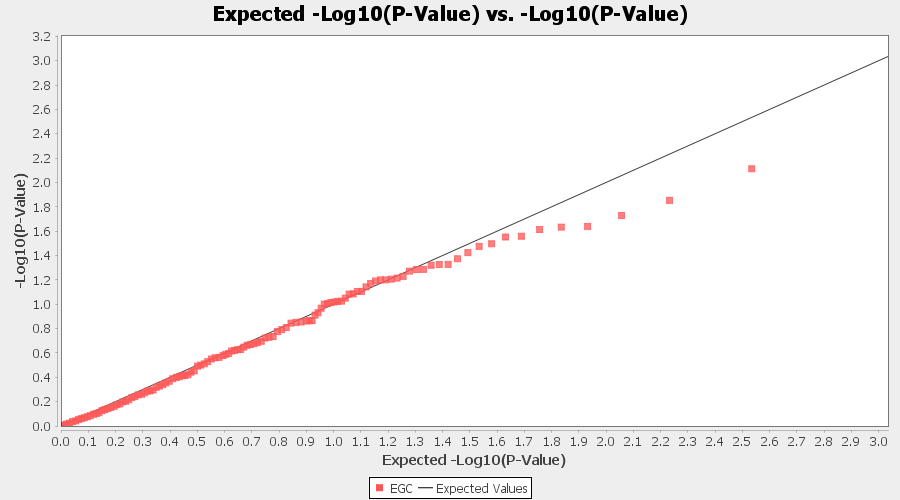

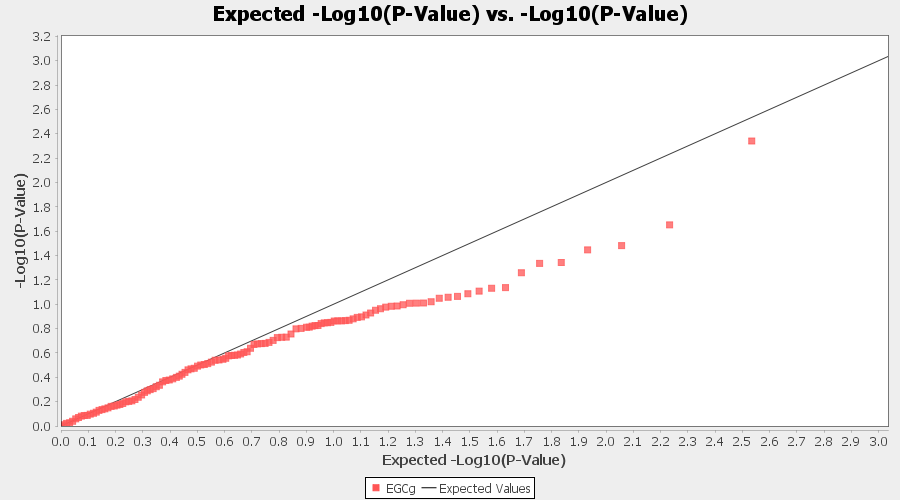

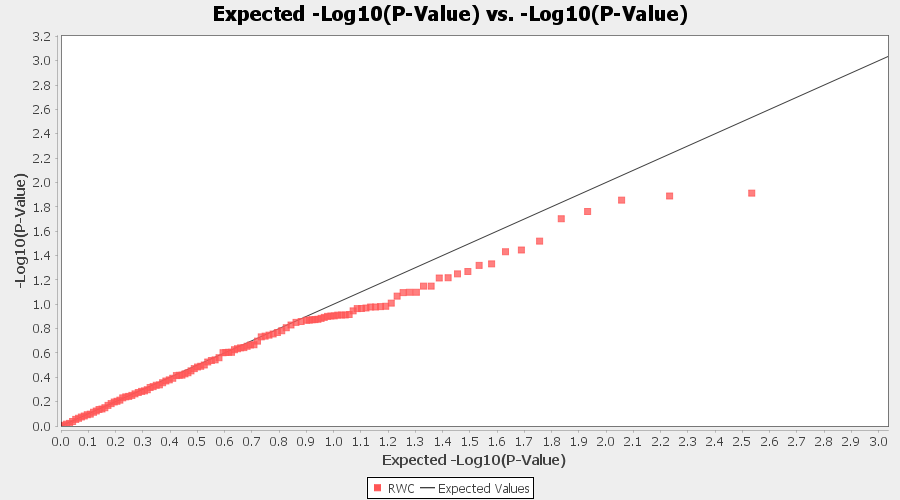

**Continued** Supplementary Figure 3. Trait-wise Q-Q association plots for mixed linear model MLM (K). ECG- Epicatechin gallate; EGC- Epicatechin gallate; EGCG- Epigallocatechin gallate; RWC- Percent relative water content

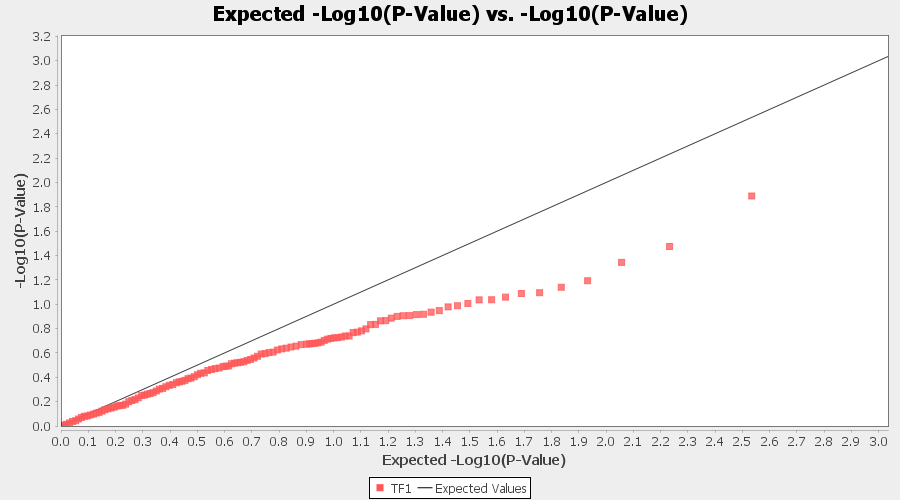

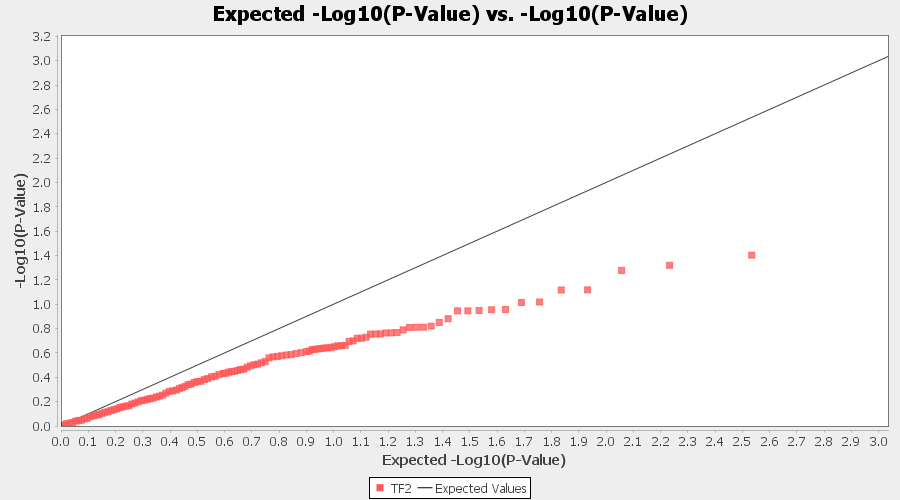

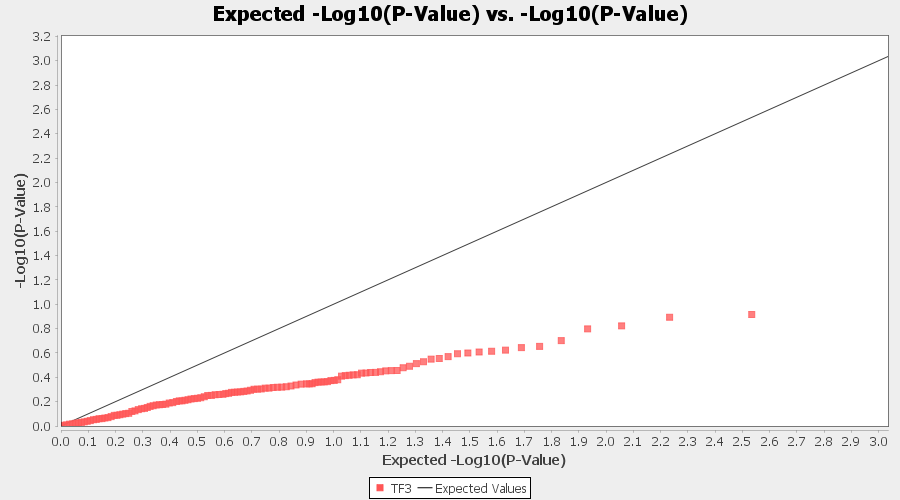

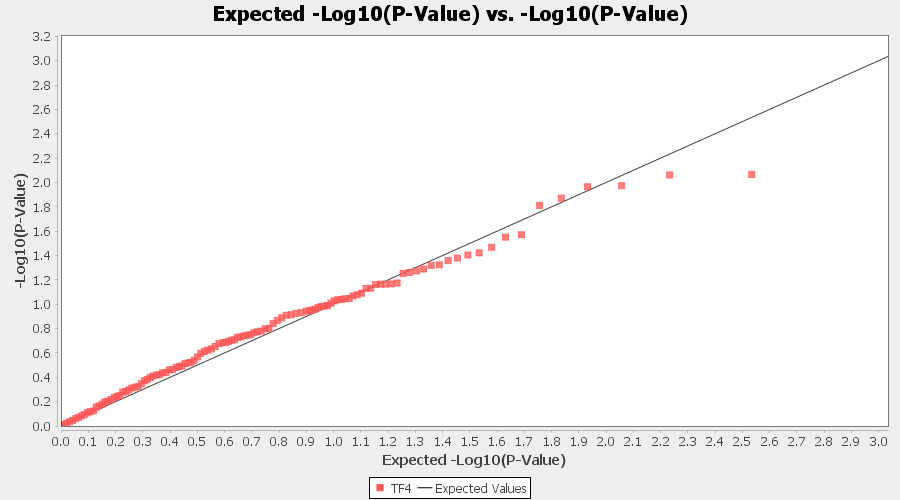

**Continued** Supplementary Figure 3. Trait-wise Q-Q association plots for mixed linear model MLM (K). TF1- Theaflavin; TF2- Theaflavin-3-gallate; TF3- Theaflavin-3’-gallate; TF4- Theaflavin-3,3’-digallate

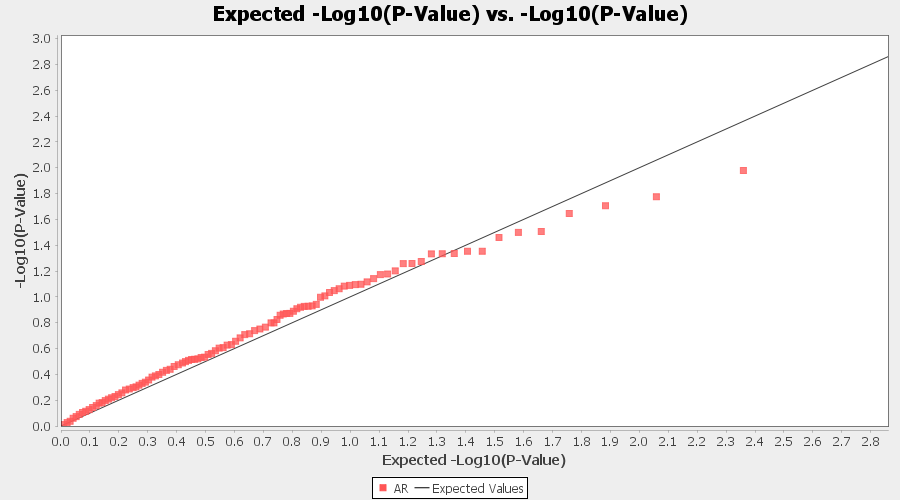

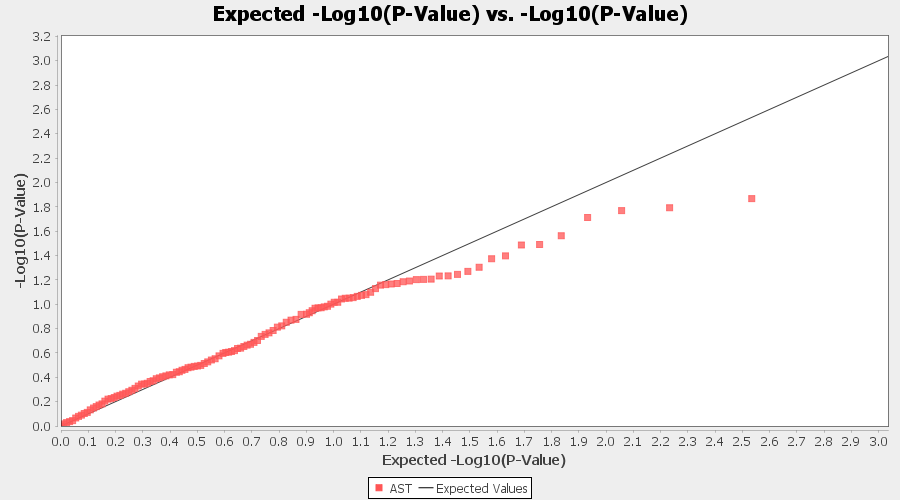

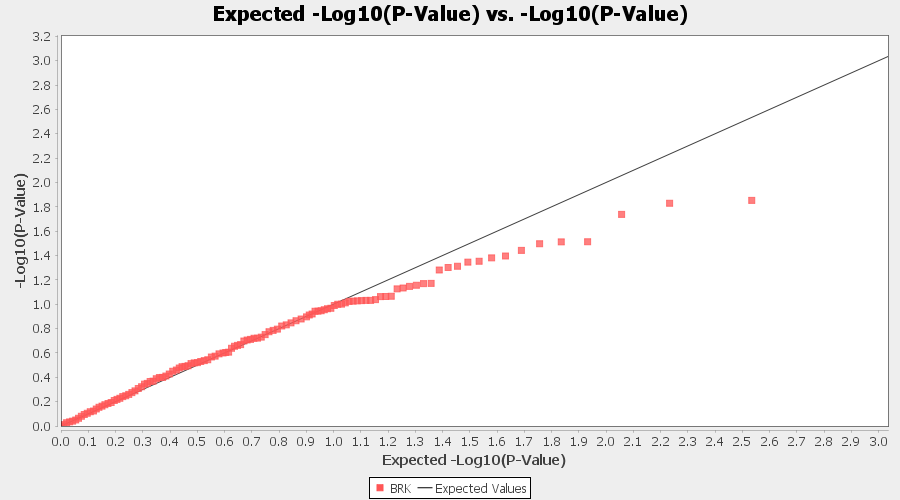

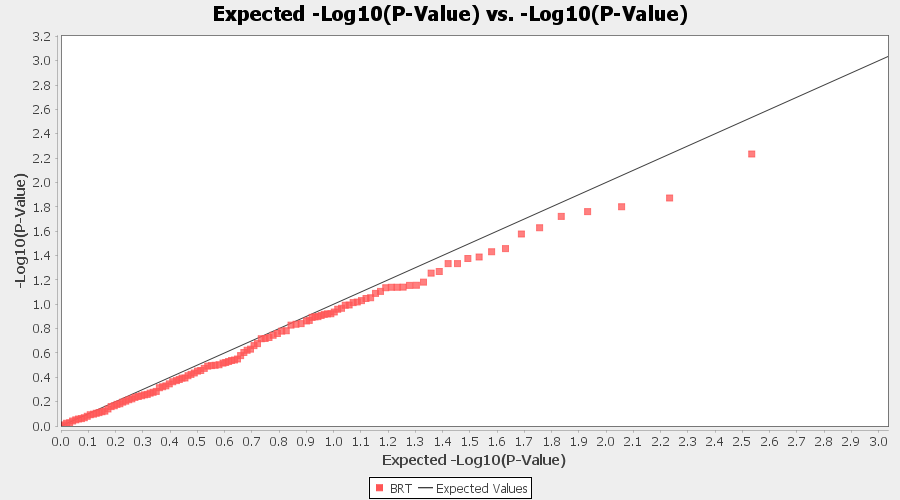

Supplementary Figure 4. Trait-wise Q-Q association plots for mixed linear model MLM (Q+K). AR- Aroma; AST- Astringency; BRK- Briskness; BRT- Brightness

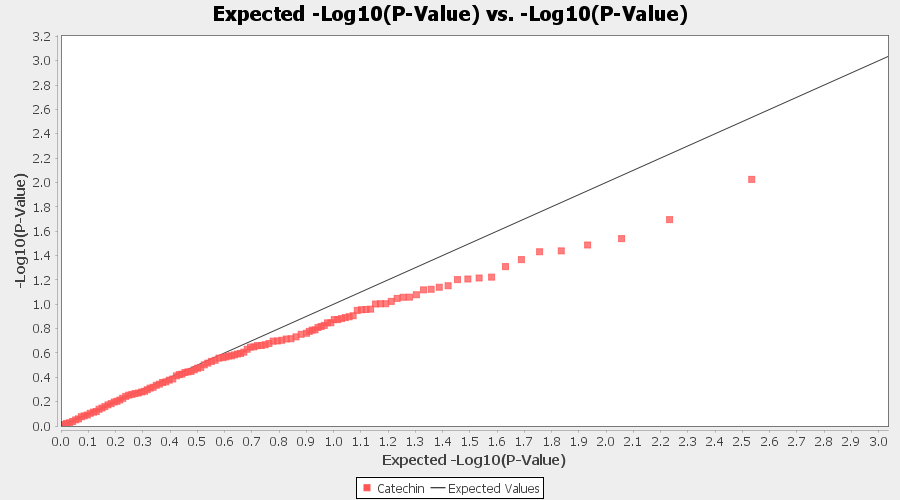

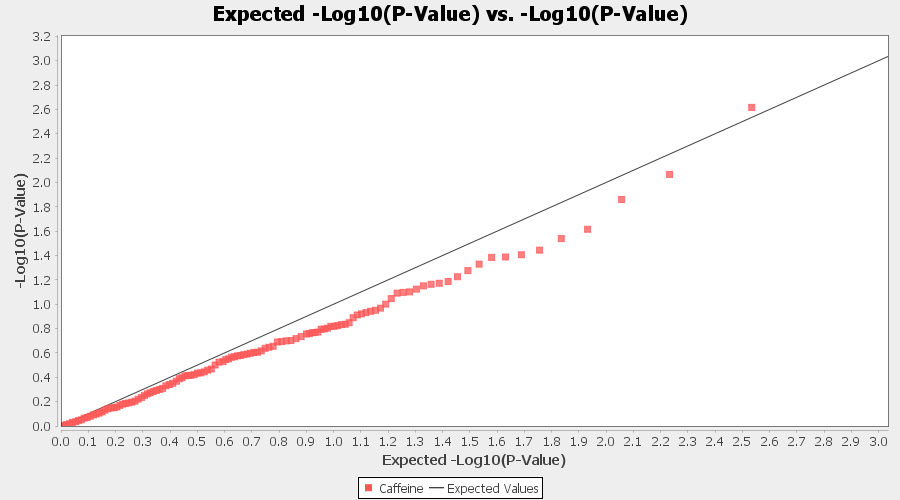

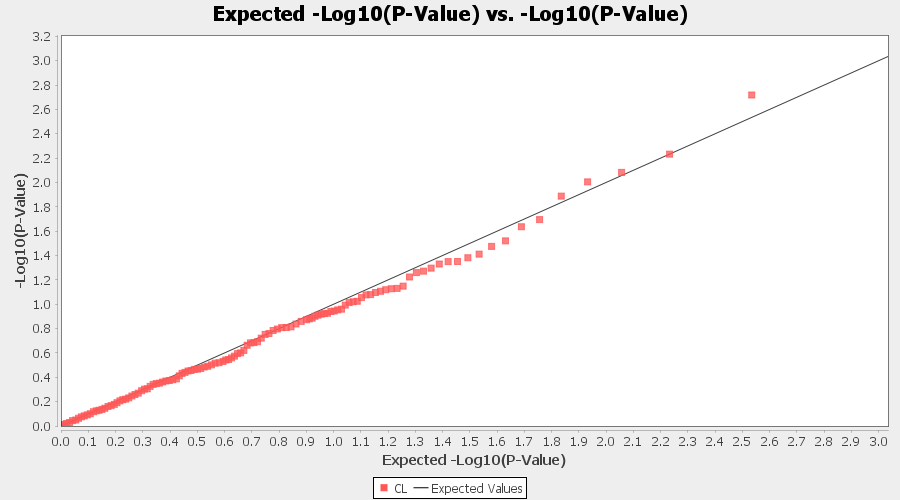

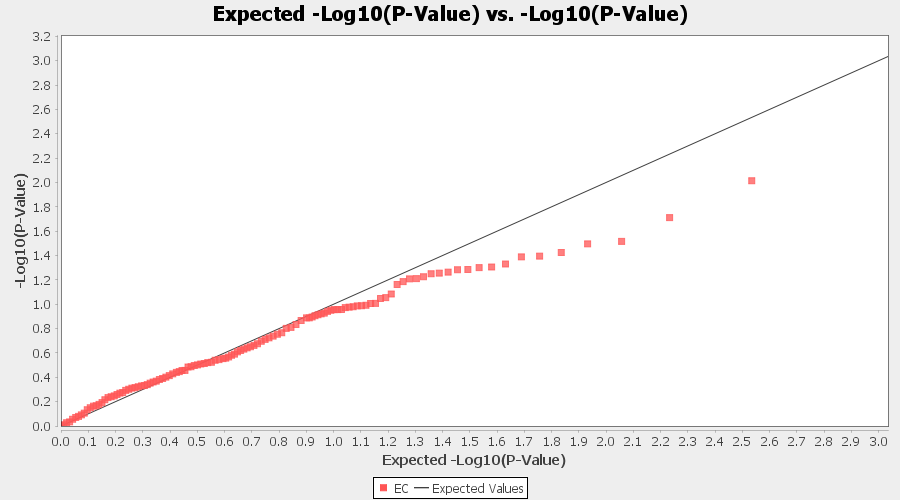

**Continued** Supplementary Figure 4. Trait-wise Q-Q association plots for mixed linear model MLM (Q+K). C- Catechin; CAFF- Caffeine; CL- Colour; EC- Epicatechin

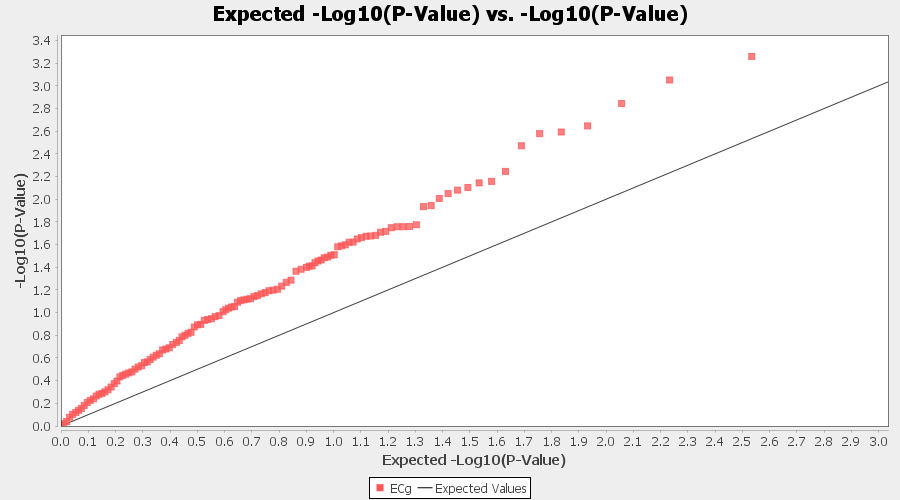

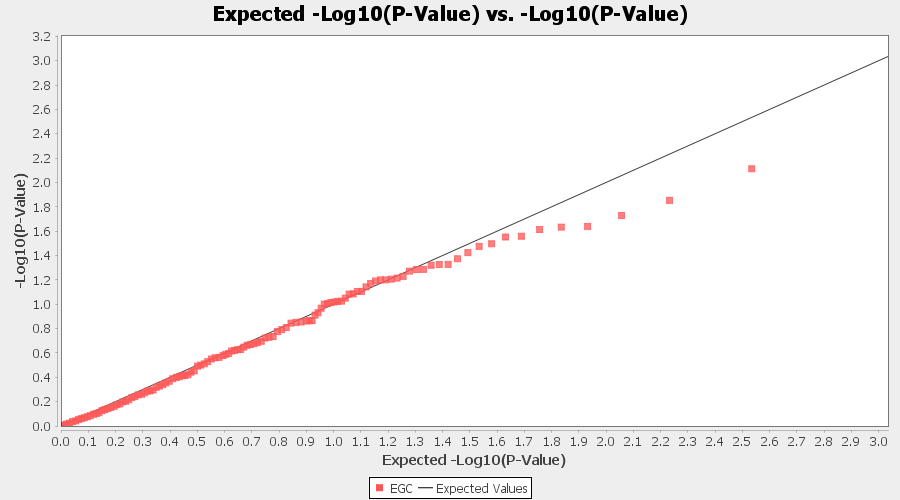

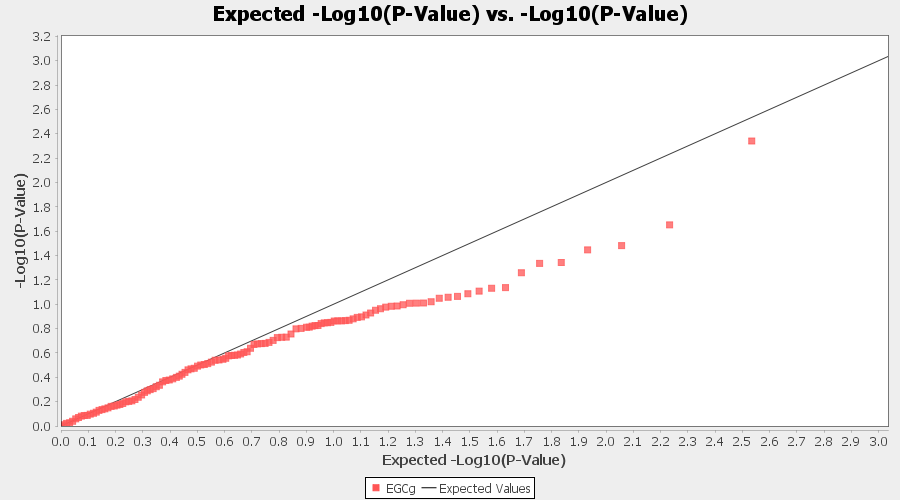

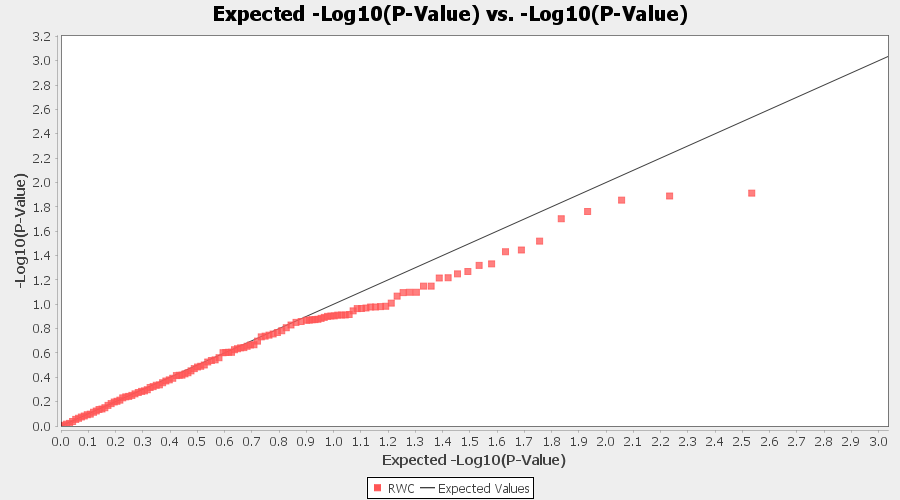

**Continued** Supplementary Figure 4. Trait-wise Q-Q association plots for mixed linear model MLM (Q+K). ECG- Epicatechin gallate; EGC- Epicatechin gallate; EGCG- Epigallocatechin gallate; RWC- Percent relative water content

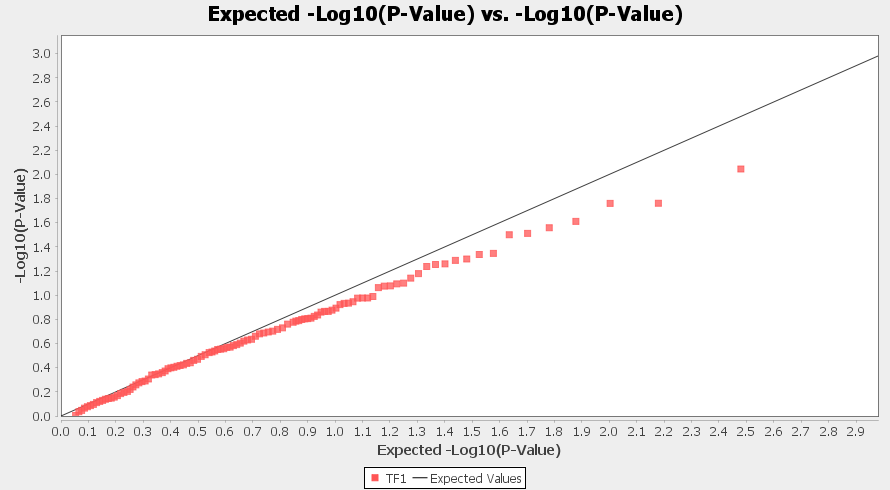

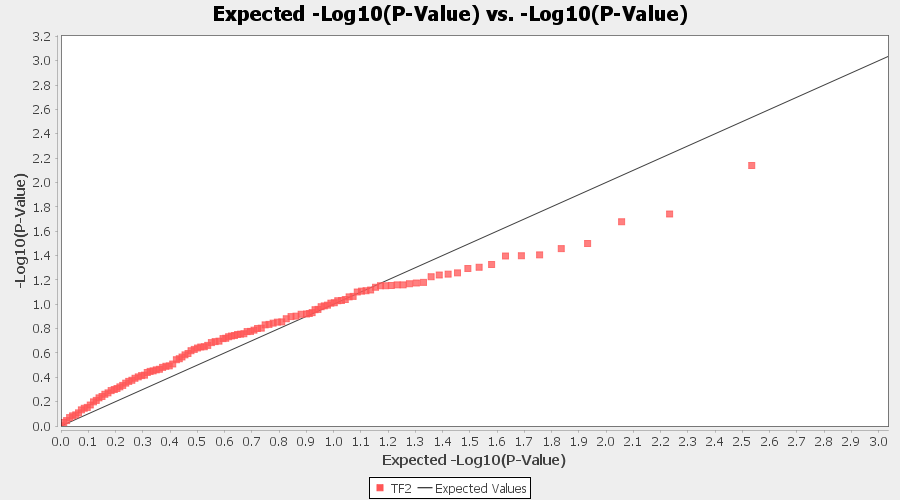

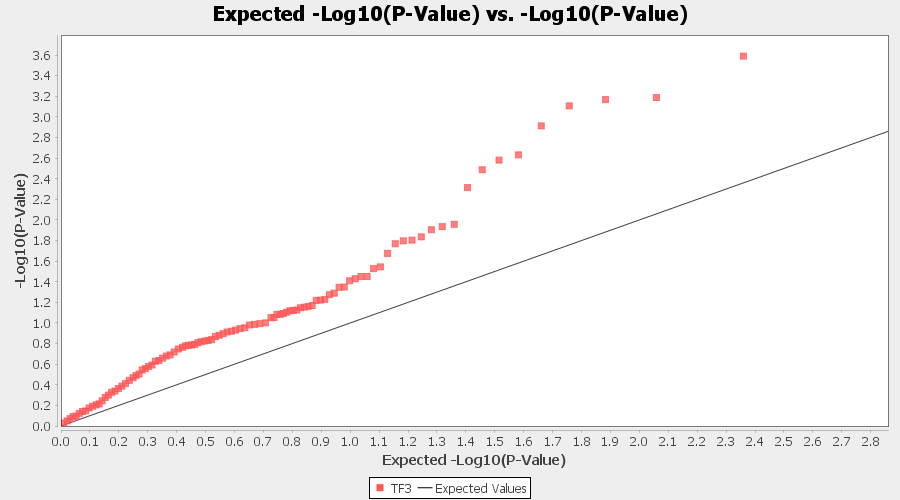

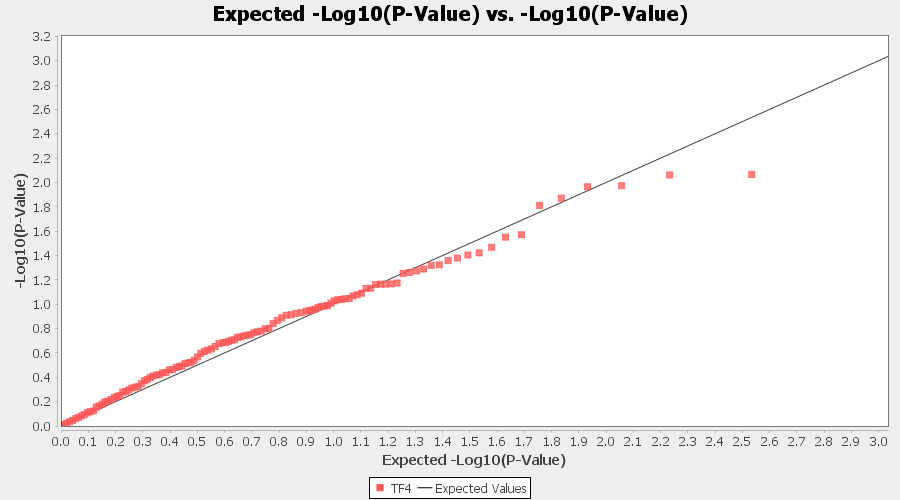
**Continued** Supplementary Figure 4. Trait-wise Q-Q association plots for mixed linear model MLM (Q+K). TF1- Theaflavin; TF2- Theaflavin-3-gallate; TF3- Theaflavin-3’-gallate; TF4- Theaflavin-3,3’-digallate
